# New glucose-abstaining *Chlorella* algae doubles mammalian cell culture longevity, boosts performance and drops serum needs enabling scaled applications

**DOI:** 10.1101/2025.05.21.655407

**Authors:** Melanie Oey, Ute Marx, Horst Joachim Schirra, James E.B. Curson, Maite Amado, Ian L. Ross, Matthew J. Sweet, Mark A. T. Blaskovich, Robert G. Parton, Ben Hankamer, Harriet P. Lo

**Affiliations:** Institute for Molecular Bioscience, The University of Queensland, Brisbane, Australia; Pforzheim University of Applied Science, Pforzheim, Germany; Griffith University, Brisbane, Australia; Centre for Advanced Imaging, The University of Queensland, Brisbane, Australia

**Keywords:** Microalgae, Co-cultivation, Tissue culture, *Chlorella*, Oxygen, Cell-waste

## Abstract

Mammalian cell culture technologies are crucial for recombinant protein production, organoid generation, medical applications, and the generation of *in vitro* cultivated meat. However, they are limited by high costs, vascular O_2_ provision, and the resultant inhibition of 3D tissue formation. Effective media usage along with oxygenation and waste management to extend culture health and longevity are key to improving all three. Microalgae, utilizing organic or inorganic CO_2_, produce O_2_ from light which complements oxygen-consuming and CO_2_-respiring mammalian cells and tissue culture. However, common microalgal cultivation conditions differ in temperature and salinity from mammalian cell cultivation environments, making co-cultivation short-lived and challenging. We screened several different microalgae species to identify locally isolated *Chlorella BDH-1* as candidate that has high growth rates in mammalian culture conditions while, unlike other *Chlorella* species, does not compete for glucose as an energy source. In mammalian cell co-culture, *BDH1* reduces cellular waste products, stabilizes pH, doubles culture longevity, increases growth performance up to 80%, and reduces expensive and ethically challenging foetal bovine serum requirements. *Chlorella BDH-1* was also non-inflammatory and tolerant of clinical antibiotics. Collectively, mammalian cell/*BDH1* co-cultivation improves tissue culture health and reduces costs, paving the path for applications in the biotechnology and medical sectors.

## 1. INTRODUCTION

### 1.1. Importance of mammalian cell culture

Mammalian cell culture is a crucial research technology with applications ranging from the study and genetically manipulation of cells and tissue organoids, to biotechnology and medical applications such as production of pesticides, pharmaceuticals (e.g. cytokines, growth factors, hormones, enzymes), vaccines, monoclonal antibodies, and products to advance gene therapy^(1)^. Meanwhile, 3D tissue culture to produce patient-specific cell-sheets for grafts has become a standard technique^(2)^. The more recent development of organoids as powerful tools to model highly complex and dynamic *in vivo* environments holds great promise for applications in various fields from drug discovery and testing to being used for regenerative medicine^(3)^. Finally, cultivated meat as early-stage technological has gained attention as an emerging field of cellular agriculture, seeking to deliver products that are traditionally made through livestock rearing in novel forms that require no, or significantly reduced, animal involvement.

Irrespective of these specific applications, key challenges centre around the culture media, which not only provides nutrients but, in a non-vascularized *in vivo* environment, also supports oxygen provision and waste management. Addressing these challenges, while optimizing cell culture procedures for scale-up, comes with significant costs in the form of a) time and labour as cell cultures usually require media changes every 2-3 days, and b) media components (e.g. amino acids and foetal bovine serum (FBS)), with the latter also introducing additional ethical concerns.

### 1.2. Limitations of cell culture applications

FBS is a growth supplement widely used in cell culture providing essential media components such as hormones, vitamins and growth factors required for cell adhesion, proliferation and metabolism^(4)^. In addition, FBS helps to improve media pH buffering capacity and to reduce damage caused by physical manipulation and stirring^(5)^. However, FBS is also the most costly component of cell culture media (∼US$600 per 500 mL bottle), with batch-to-batch variability in quality and composition, and is associated with significant ethical concerns^(5)^. The potential for bacterial or viral contamination in FBS is another important consideration for clinical and cultivated meat applications. Therefore, significant work has been conducted to reduce the need for FBS in cell culture media. While there are ∼260 unique mammalian cell types that can be cultured in serum-free media, these are typically even more expensive due to their complex formulations and the necessity to include recombinant growth factors replacing FBS. There is currently no single serum-free media alternative that can replace FBS to culture all cell types^(5)^.

### 1.3. Microalgal characteristics required to complement cell culture systems

Microalgae are photosynthetic single-cellular organisms, which can grow by using organic or inorganic carbon sources (CO_2)_ and light to produce oxygen. Thus, several microalgae species have been tested to complement mammalian cell culture systems, to provide O_2_ to and absorb CO_2_ from mammalian cells. However, microalgal survival in mammalian cell culture conditions has proved challenging^(6-8)^, and these challenges must be considered when selecting the most suitable microalgae for the task.

#### 1.3.1. Temperature

One challenge of co-cultivating microalgae with mammalian cells is the difference in their preferred growth temperature. Microalgae are usually grown at a temperature range of 15– 30 °C, with an optimum temperature of 20–25 °C^(9, 10)^, whereas mammalian cells require higher temperatures due to their adaptation to the core body temp (∼37 °C in humans and 36.6 °C in mice^(11)^), and are often less tolerant to temperature fluctuations.

#### 1.3.2. Salinity

Mammalian blood and tissue culture medium typically contains ∼150 mM NaCl^(12)^. In comparison, freshwater has less than ∼17 mM total salinity, and seawater in tropical areas can have salinities equivalent to ∼600 mM NaCl or more^(13)^. Freshwater strains such as the model microalgae *Chlamydomonas reinhardtii* can be salt-adapted, but adaption often does not yield optimal growth. For example, in *C. reinhardtii* 150 mM NaCl stress leads to flagellar resorption, reduction in cell size, slower growth rates and aggregation^(14)^. Only after 17 months (1255 generations) under a regime of serial dilutions every 2–3 days, did the cells yield similar growth rates to those cultivated in salt-free media. *Chlorella vulgaris* was reported to tolerate a large range of salinity, but also requires adaptation procedures to increase resistance against high salinity^(15)^. Saltwater species such as *Nannochloropsis sp.* or *Dunaliella sp.* are often cited as microalgal candidates for desalination, indicating adaptability to different salinities^(16)^. However, the salt content of mammalian cell media (150 mM NaCl) may not be sufficiently saline to support the growth of these or other marine strains. For example, *Nannochloropsis* and *Dunaliella* are reported to grow best in salinities above 375 mM NaCl^(17, 18)^. In summary, strains capable of growth in brackish water are likely to be most closely adapted to the salinity levels required by mammalian cells.

#### 1.3.3. Carbon source

While temperatures and salinities are already a challenge for algal growth, in co-culture the maintenance of optimum growth conditions and nutrients for both, algal and mammalian cells, must be achieved. It is thus important to select microalgae that do not compete with the mammalian cells for nutrients such as glucose, as this serves as an energy source for mammalian cells. We have previously reported that a large number of microalgae (∼50% of ∼100 tested microalgae strains) were capable of utilizing glucose as carbon source^(19, 20)^.

In summary, the ideal microalgal species to support mammalian cells in co-culture tolerates the same temperature, salinity, and composition of the mammalian culture medium, while providing oxygen, reducing CO_2_ and fermentative waste products, and not competing for nutrients, especially glucose.

### 1.4. Our study

Here we present the microalgal strain *Chlorella BDH-1* which meets the above requirements. Out of 10 different strains comprising 5 different microalgae species we identified an Australian microalgal strain, *Chlorella BDH-1,* which tolerates mammalian cell culture conditions while simultaneously sparing glucose, the primary mammalian cell energy source. We chose the well-characterized C2C12 mouse myoblast cell line for co-cultivation, as muscle cells are among the most metabolically active cells in the body^(21)^, thus serve as a proof-of-concept model for the cultivated muscle-meat industry^(22)^.

## 2. RESULTS and DISCUSSION

### 2.1. Microalgae and mammalian cell selection for co-cultivation

Microalgae are single celled phototrophs that are adapted to diverse planetary habitats. We therefore hypothesized that there would be an ideal microalga compatible with the cell culture conditions required for mammalian cells. After careful consideration, we chose 10 different promising algae strains to be tested for suitability in co-cultivation with mammalian cells. This included freshwater and saltwater microalgae from 5 different species (Table 1).

**Table 1:**
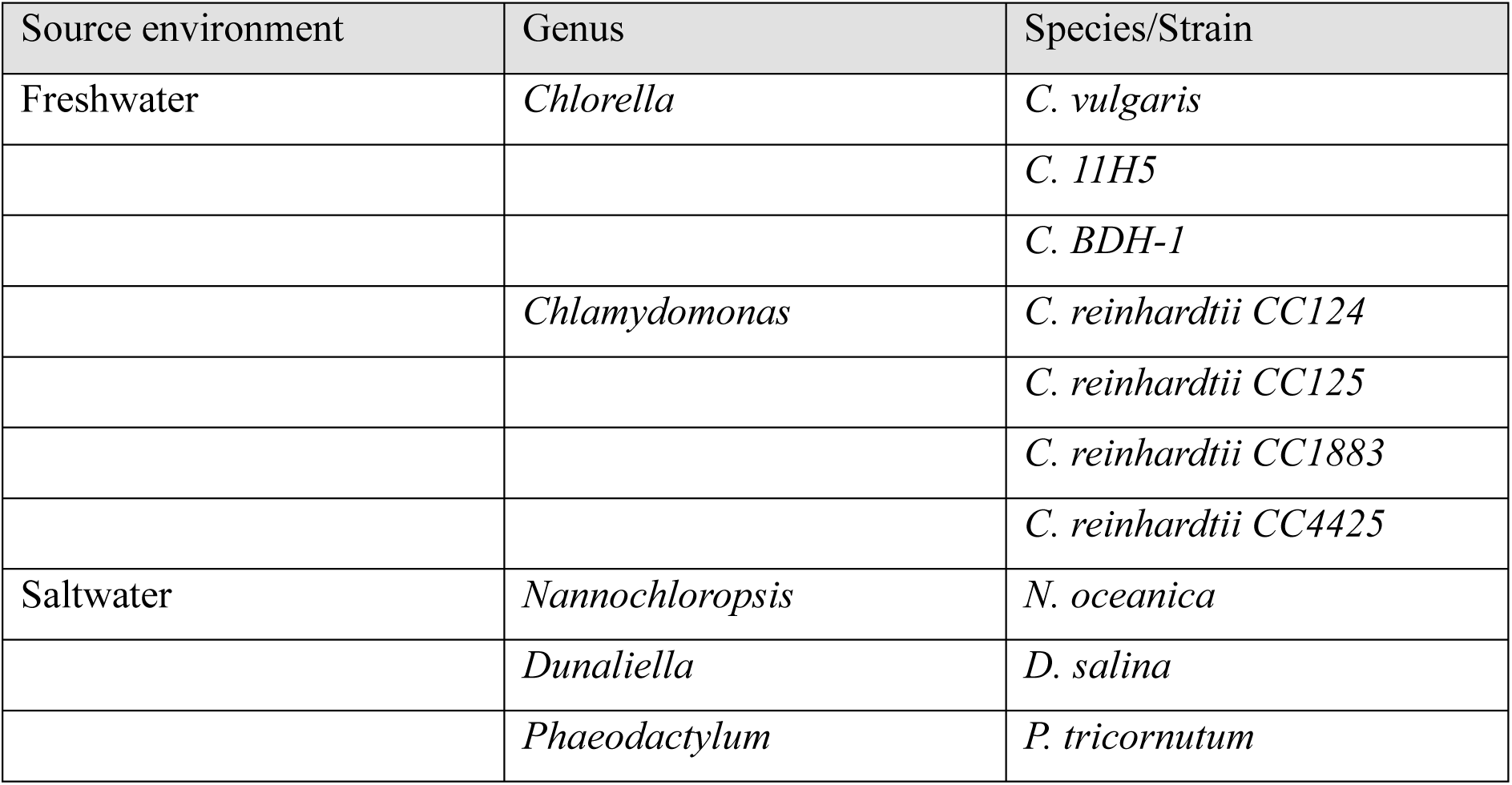
Microalgae species and strains tested in this study.

| Source environment | Genus | Species/Strain |
| --- | --- | --- |
| Freshwater | *Chlorella* | *C. vulgaris* |
|  |  | *C. 11H5* |
|  |  | *C. BDH-1* |
|  | *Chlamydomonas* | *C. reinhardtii* CC124 |
|  |  | *C. reinhardtii* CC125 |
|  |  | *C. reinhardtii* CC1883 |
|  |  | *C. reinhardtii* CC4425 |
| Saltwater | *Nannochloropsis* | *N. oceanica* |
|  | *Dunaliella* | *D. salina* |
|  | *Phaeodactylum* | *P. tricornutum* |

First, we established a growth performance baseline for each microalgae strain at different temperatures and in different media.

Microalgae were cultured under illumination (Fig. 1A) at ambient room temperature (RT) or in a cell culture incubator in three different standard algae cultivation media: (1) Freshwater TAP medium^(23)^, which provides acetate as a carbon source, (2) Photoautotrophic PCM medium^(24)^, and (3) photoautotrophic saltwater F/2 medium ^(25)^), as well as standard cell culture medium. Microalgal chlorophyll, with an absorbance peak at 680 nm is a useful indicator of cell viability and growth^(26)^, and thus was used to measure microalgal growth (Fig. 1B-D). At RT (Fig. 1BC), all tested freshwater strains grew well in TAP medium. The best growth under photoautotrophic conditions in PCM medium was displayed by *C. reinhardtii* CC125 and the local isolate *C. BDH-1*. As expected, the three saltwater strains (*N. oceanica*, *D. salina* and *P. tricornutum*) grew poorly in TAP and PCM medium. In saltwater F/2 medium *N. oceanica* showed the best growth rates. Surprisingly, the local strains (*C. 11H5 and C. BDH-1)* showed comparable growth in F/2 medium to the saltwater strain *N. oceanica*, without prior salt adaption. As expected, the freshwater *Chlamydomonas* strains did not show any growth in F/2 medium.

**Figure 1:**
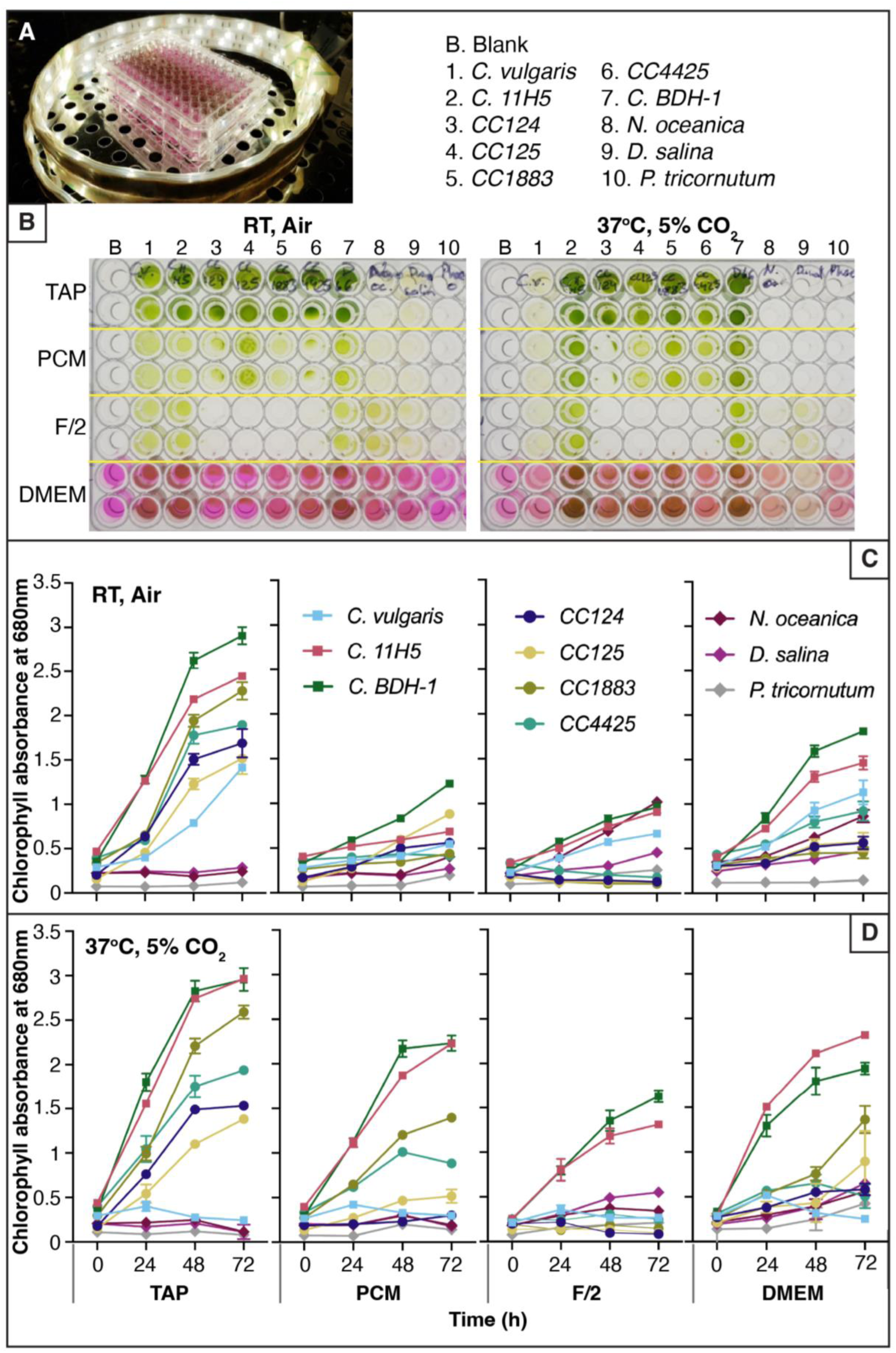
Cultivation of 10 different microalgae strains in different media. **(A)** Light setup providing 100 µmol photons m^-2^ s^-1^ constant illumination to 96-well plates. **(B)** Plates imaged after 72 h at ambient RT (25°C, 0.04% CO2) or in a cell culture incubator (37°C, 5% CO_2_). **(C-D)** Microalgae growth measured as chlorophyll absorbance at 680 nm (mean±SD, n=3) at RT (C) or 37 °C (D) in TAP, PCM, F/2, or DMEM growth media over the course of 72 h in 96 well-plates.

To assess growth under conditions compatible with mammalian cell culture, microalgae were next grown in different algae media in a standard cell culture incubator at 37°C (Fig. 1B and 1D). Notably, all freshwater strains except *C. vulgaris* grew well in TAP medium with the Australian isolates *C. 11H5* and *C. BDH-1* performing best. The relatively poor performance of freshwater strain *C. vulgaris* was attributed to its optimum growth temperature being ∼25°C^(27)^. In photoautotrophic PCM medium, *C. 11H5* and *C. BDH-1* also performed best, followed by the *Chlamydomonas* strains CC1883 and CC4425. As expected, the saltwater strains did not show any growth in TAP or PCM media. Interestingly, in F/2 saltwater medium, *C. 11H5* and *C. BDH-1* showed higher growth compared to all saltwater strains, which grew poorly at 37°C.

The ability of the microalgae strains to grow in standard cell culture medium (DMEM) was also tested at RT and 37°C (Fig. 1B-D). As observed above, *C. vulgaris* growth was abolished at 37°C. Increased growth of *C. 11H5* and *C. BDH-1* was observed in DMEM at 37°C compared to RT, although the growth was not as high as in TAP media. The higher photoautotrophic growth at 37°C for *C. 11H5* and *C. BDH-1* is most likely due to the high ambient CO_2_ and temperature tolerance. The major organic carbon source in DMEM medium is glucose, which serves as an energy source for mammalian cells. The genus *Chlorella* is typically reported to utilize glucose as a carbon source ^(20, 28)^. However, the reduced growth in DMEM compared to TAP, as well as the similar level of growth in photoautotrophic conditions (PCM) suggest that glucose does not effectively substitute for acetate as a carbon source for *C. 11H5* and *C. BDH-1*. Consequently, they would not be expected to compete with mammalian cells for glucose.

### 2.2. Species identification of locally isolated *BDH-1* strain

To identify the precise species of the locally isolated *BDH-1* strain, genomic DNA was isolated and two sections of the 18S ribosomal RNA gene were amplified and sequenced. A BLAST search^(29)^ was conducted against the obtained sequences, and the microalgal species was identified as a close relative to *Chlorella sorokiniana* with 99.4% homology to NCBI Accession KT852969.1. After the identification as *Chlorella*, 16S ribosomal DNA was amplified using universal as well as *Chlorella-*specific primers^(30)^; sequencing alignments confirmed C. *BDH-1* to indeed be closely related to *Chlorella sorokiniana* (99% sequence similarity to *C. sorokiniana* NCBI Accession KJ742376.1). *Chlorella sorokiniana* is a well-known and characterized microalgae species, ‘Generally Recognized As Safe” by the FDA and approved for food consumption (GRN986). Due to its performance in the above-described growth experiments and its close relation to *C. sorokiniana*, we chose *Chlorella BDH-1* over *Chlorella 11H5* as our best candidate for subsequent experiments.

### 2.3. *Chlorella BDH-1* cultivation in C2C12 spent media

As media composition changes during cell cultivation, shifting from nutrient abundance to waste product accumulation, we next investigated *C. BDH-1* growth performance and duration at 37 °C in spent DMEM media at atmospheric CO_2_ levels (Fig. 2A and B). Growth in *fresh DMEM* media was again comparable to photoautotrophic growth, indicating that *C. BDH-1* did not utilize glucose as a carbon source for growth. These results were also confirmed by NMR below. Interestingly, in spent DMEM media the microalgae showed growth comparable to acetate-containing TAP media, indicating that a utilizable carbon source had been released into the media by the C2C12 cells. Visual assessment and growth measurement of chlorophyll absorbance at 680 nm (Fig. 2A and B) did not show a decline in chlorophyll content, indicating that the cells were still healthy and viable after 96 hours.

**Figure 2:**
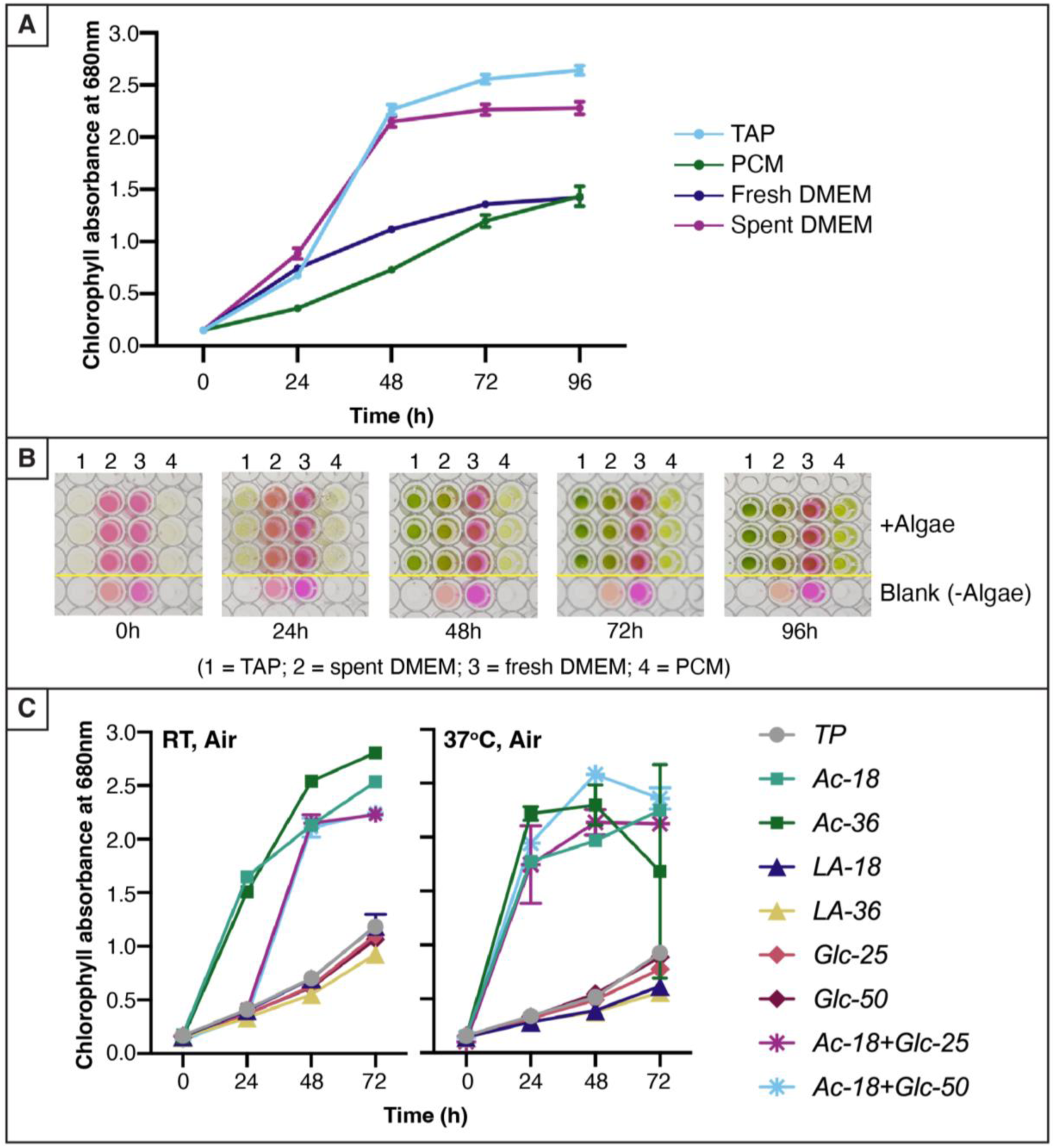
*Chlorella* BDH-1 growth in spent cell-culture media and different organic carbon sources. Growth was determined via chlorophyll absorbance at 680 nm. **(A)** Microalgal growth at 37°C in TAP media, PCM media, and mammalian DMEM growth media before (fresh DMEM) or after (spent DMEM) 72 h cultivation with C2C12 cells (mean±SD, n=3). **(B)** Images of 96-well plates for samples in (A). **(C)** C. BDH-1 growth in TP media providing different organic carbon sources at room temperature (left) and 37°C (right) (mean±SD, n=3). TP media was supplemented with 18 mM (LA-18) or 36 mM (LA-36) lactate, 25 mM (Glc-25) or 50 mM (Glc-50) glucose (Glc-25 and Glc-50, respectively), 18 mM (Ac-18, similar to TAP media) or 36 mM (Ac-36) acetate, or 18 mM acetate with 25 mM or 50 mM (Ac18+Glc50) glucose, (Ac18+Glc25 or Ac18+Glc50, respectively).

### 2.4. Utilization of acetate, lactate and glucose by *BDH-1* at different temperatures

Glucose, acetate, and lactate are organic carbon compounds that influence the performance of mammalian cell culture, either by being used as nutrients for cell growth or excreted as metabolic waste compounds into the media and thus are likely candidates for the released carbon source. To assess the effect of these organic carbon sources on *C. BDH-1*, we tested microalgal growth in commonly used microalgae TP media^(23, 24)^ supplemented with various concentrations of acetate, glucose or lactate (Fig. 2). Growth was then monitored using chlorophyll absorption at 680 nm at RT and 37°C both at atmospheric CO_2_ to eliminate additional effects of supplemented CO_2_.

Similar to previous findings in media supplemented with acetate (TAP), *BDH-1* showed faster growth onset at 37 °C than at RT (Fig. 2) after 24 h, which was much more pronounced in higher acetate concentrations, although final chlorophyll absorption at 48 h was comparable. In cultures provided with both acetate and glucose, no growth difference was observed between ± glucose supplementation, indicating that *C. BDH-1* preferred acetate over glucose as a carbon source. At 72 h and 37°C, all cultures that were provided with acetate showed a decline in chlorophyll absorbance, presumably due to the fast growth onset and subsequent plateau phase caused by depletion of other nutrients. We also noted that at both temperature conditions *C. BDH-1* grew more poorly in media containing glucose or lactate as the sole carbon source compared to acetate. Growth in media containing glucose and lactate was comparable to samples without any organic carbon supply (TP) (Fig. 2). We therefore conclude that *C. BDH-1* does not effectively utilize glucose or lactate for growth and is unlikely to compete with mammalian cells for glucose as a carbon source.

### 2.5. Optimal ratio of co-cultivated microalgae to C2C12 cells supports full usage of cell culture media

We next aimed to assess the best algae-to-mammalian cell ratio (Fig. 3A) to improve mammalian cell growth. Although preliminary experiments showed that both cell types could be co-cultivated, we utilized transwell inserts (Fig. 3B) to separate microalgae from C2C12 cells to simplify viability assays but allow media diffusion. To identify the best ratio, a set number of *C. BDH-1* microalgae cells were applied to 20,000 C2C12 cells, and mammalian cell growth was observed over 6 days. The algae-containing inserts were removed to assess the number of metabolically active, viable C2C12 cells using resazurin-based assays. After 2 days of co-cultivation with microalgae, we consistently observed similar or higher numbers of metabolically active mammalian cells compared to the no-algae control (Fig. 3A, Supplementary Table ST1). However, we determined the optimum C2C12 to *C. BDH-1* microalgae cell ratio to be 1:20, yielding up to 1.8-fold more metabolically active C2C12 cells compared to the algae-free control.

**Figure 3:**
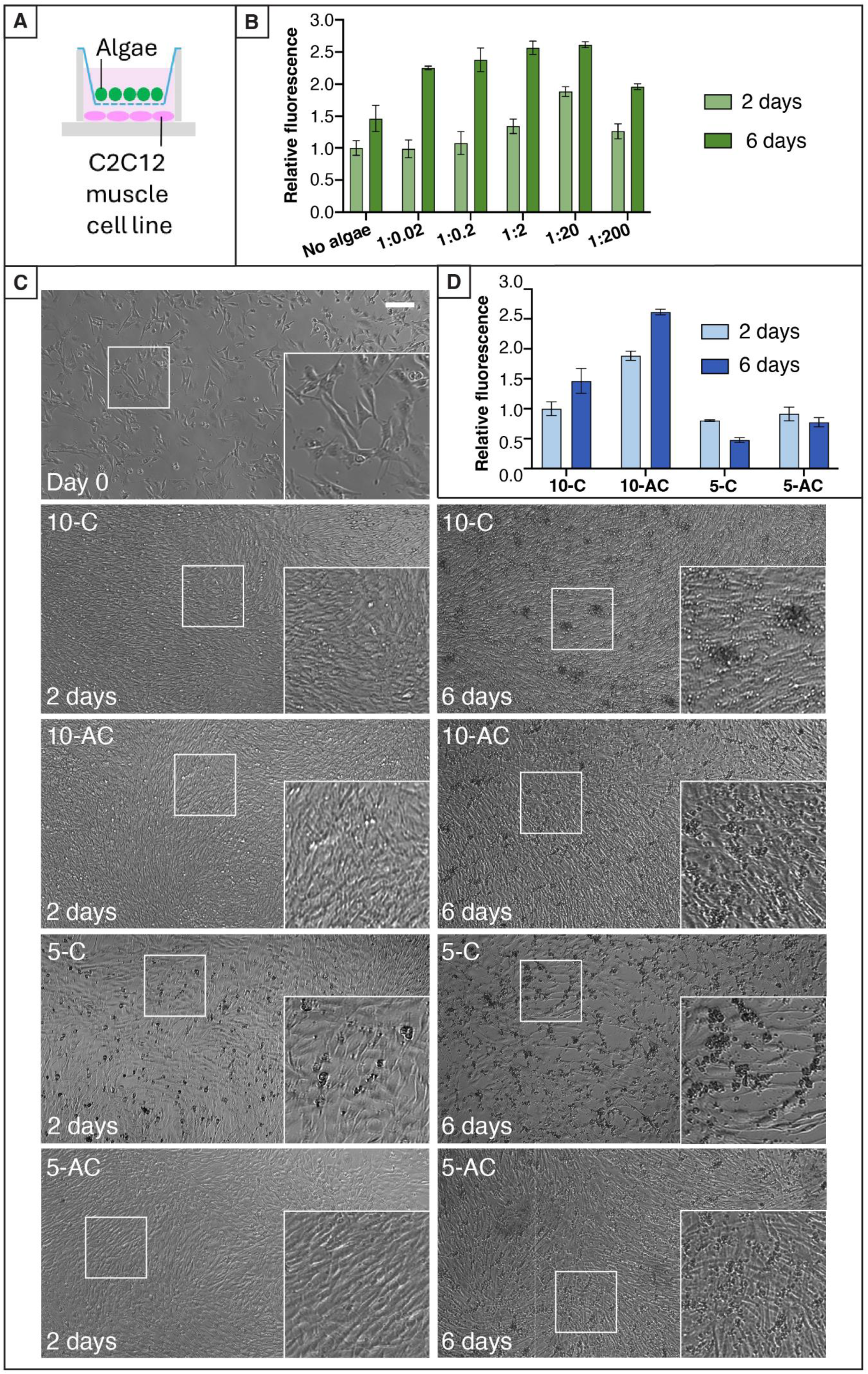
Optimum co-culture cell ratios for extended cell culture longevity and reduced serum requirements. **(A)** Schematic cell separation using transwell inserts. **(B)** Optimization of C2C12:*C. BDH-1* cell number ratio after 2 or 6 days co-cultivation assessed via resazurin-based assays. Quantitation relative to “no algae” control after 2 days cultivation. **(C)** Representative brightfield images of C2C12 cells 24 h after plating (day 0), and after 2 or 6 days cultivation in standard DMEM media with 5% or 10% FBS, with or without microalgae co-cultivation. Insets represent boxed areas. Bar, 200 µm. Replicates see Fig. S1. **(D)** Average metabolic activity of C2C12 cells after 2 or 6 days cultivation in standard DMEM media with 10% FBS or 5% FBS, with or without C. BDH-1 co-cultivation. Quantitation relative to “10-C” control after 2 days cultivation. **(B) and (C):** Mean±SEM from up to 6 independent experiments with up to 3 technical replicates each (see individual experiments in supplementary ST1 and ST2). See Table 2 for media abbreviations.

**Table 2:**
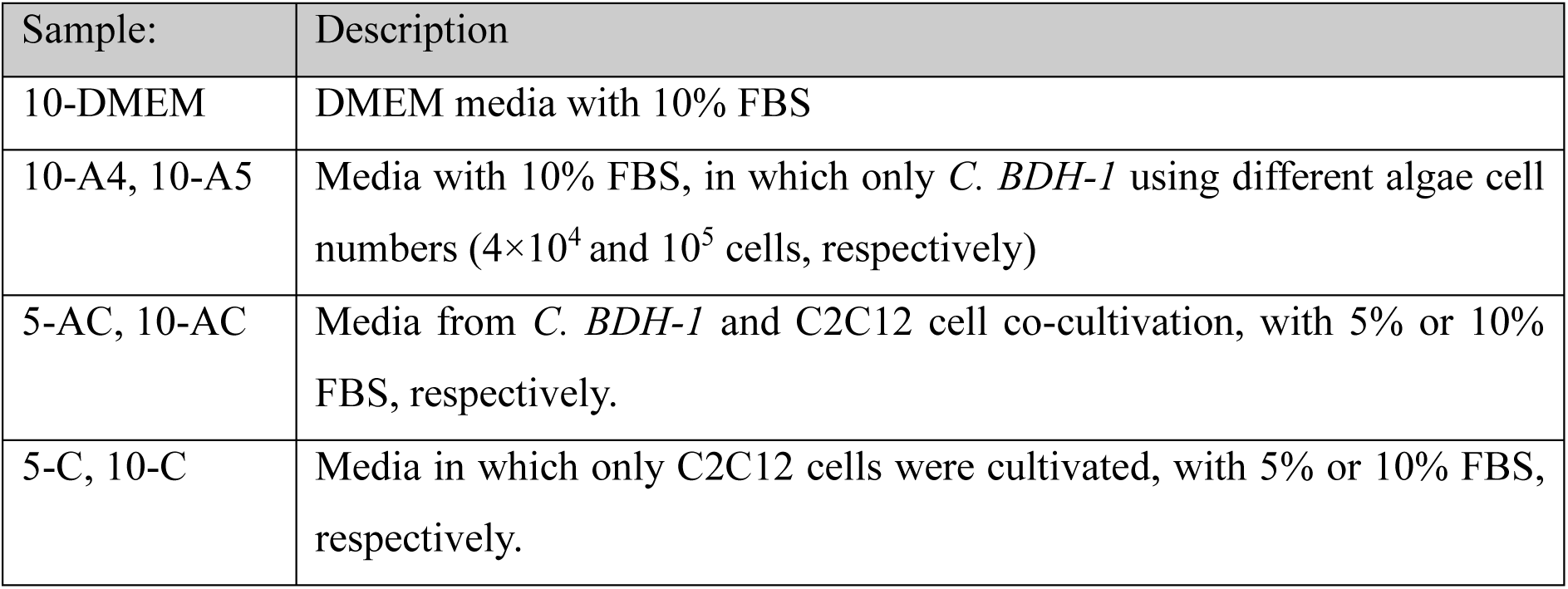
Sample abbreviations and description of the respective media.

As it is common practice to exchange culture media every 2-3 days, to replenish nutrients, we also assessed C2C12 cell viability after 6 days of co-cultivation with microalgae (without media exchange) to test the robustness of the co-cultivation system. Remarkably, the determined optimum C2C12:microalgae ratio of 1:20 cells not only supported better growth at day 2 but also supported extended C2C12 growth and viability to day 6, doubling the typical culture lifespan while yielding increased numbers of viable cells, up to 2.5-fold compared to the algae-free control (Fig. 3A). This indicates that the microalgae improve culture conditions, allowing the remaining growth components in the DMEM media to be utilized. It further suggests that the common requirement for frequent culture media exchange is more due to waste product accumulation rather than for nutrient depletion. Thus, valuable and usable components are disposed of, adding to high culture media costs.

### 2.6. Microalgae co-cultivation allows a reduction in serum requirements

Typically, FBS is one of the most expensive components in mammalian cell culture. We thus explored whether co-cultivation with microalgae could reduce the amount of FBS required. C2C12 cells were seeded into a 24-well transwell plate and incubated for 24 h. The culture medium was then replaced by medium containing 10% FBS or 5%FBS, respectively, and cultured with or without microalgae cells (at a ratio of 1:20) for 2 and 6 days. C2C12 cell growth was monitored visually (Fig. 3C), and metabolic activity was assessed using a resazurin-based viability assay (Fig. 3D). Sample abbreviations and respective cultivation media are listed in Table 2.

On day 2, C2C12 cells grown in 10% FBS with (10-AC) or without algae (10-C) as well as with algae in 5% FBS (5-AC) appeared healthy, whereas large numbers of dead cells (identified as dark spherical bodies) and reduced, disrupted confluence was observed in cells grown in 5% FBS without microalgae (5-C) (Fig. 3C). These differences were more pronounced at day 6; all C2C12 dishes co-cultivated with microalgae showed healthier and more confluent growth compared to samples cultivated with 10% FBS without microalgae (10-C). Consistent with these observations, metabolic resazurin-based viability assays revealed higher numbers of metabolically active cells at day 2 when grown in 10% FBS and co-cultivated with microalgae, in comparison to control samples grown without microalgae (10-C) (Fig. 3D and supplementary Fig. S1). Cells grown in 5% FBS with microalgae were similar to control samples. This indicates that microalgae support mammalian cell growth even in early stages (also seen in Fig. 3A). The number of metabolically active C2C12 cells in 10-AC increased over 6 days reaching up to 2.6-fold the cell number compared to algae-free 10-C culture at day 2, whereas cell viability of 10-C increased only to 1.4-fold. At day 6 co-cultivated 5-AC samples showed higher numbers of metabolically active C2C12 cells compared to the control samples 10-C at the same time point (Supplementary ST2). However, overall metabolic cell activity had decreased compared to 5% FBS samples at day 2. This indicates that microalgae can support faster cell growth with extended cultivation timeframes, while their ability to compensate for reduced serum concentrations is limited.

### 2.7. Microalgae extend tissue culture longevity by maintaining pH and decreasing waste product accumulation

#### 2.7.1. pH stabilization

Oxidative phosphorylation and anaerobic glycolysis are two means of energy production in mammalian cells, each with different glucose-to-energy conversion efficiencies, oxygen requirements and waste production (Fig. 4A). Oxidative phosphorylation is a cell’s primary and most efficient way of producing ATP, with little production of acidic fermentation byproducts such as lactate and acetate^(31)^. However, if oxidative phosphorylation is inhibited, cells switch to glycolysis to produce ATP generating considerably more lactate. An artificial cell culture system, lacking blood supply and thus oxygen provision and waste removal, is prone to waste accumulation, leading to a drop in the pH, and a reduction in the utilization of full media components. Phenol red is a pH indicator^(31)^ (Fig. 4B) that is commonly used in mammalian cell culture medium to monitor growth conditions, metabolic status and cell health, providing a quick check for culture health^(32)^. Notably, at day 6 cultivation in DMEM with 10%FBS we observed a clear colour difference between the co-cultivation wells with algae (+ A, Fig. 4C), which appeared pinker. In comparison, wells that only contained C2C12 cells (- A, Fig. 4C) appeared more yellow indicating a lowered pH.

**Figure 4:**
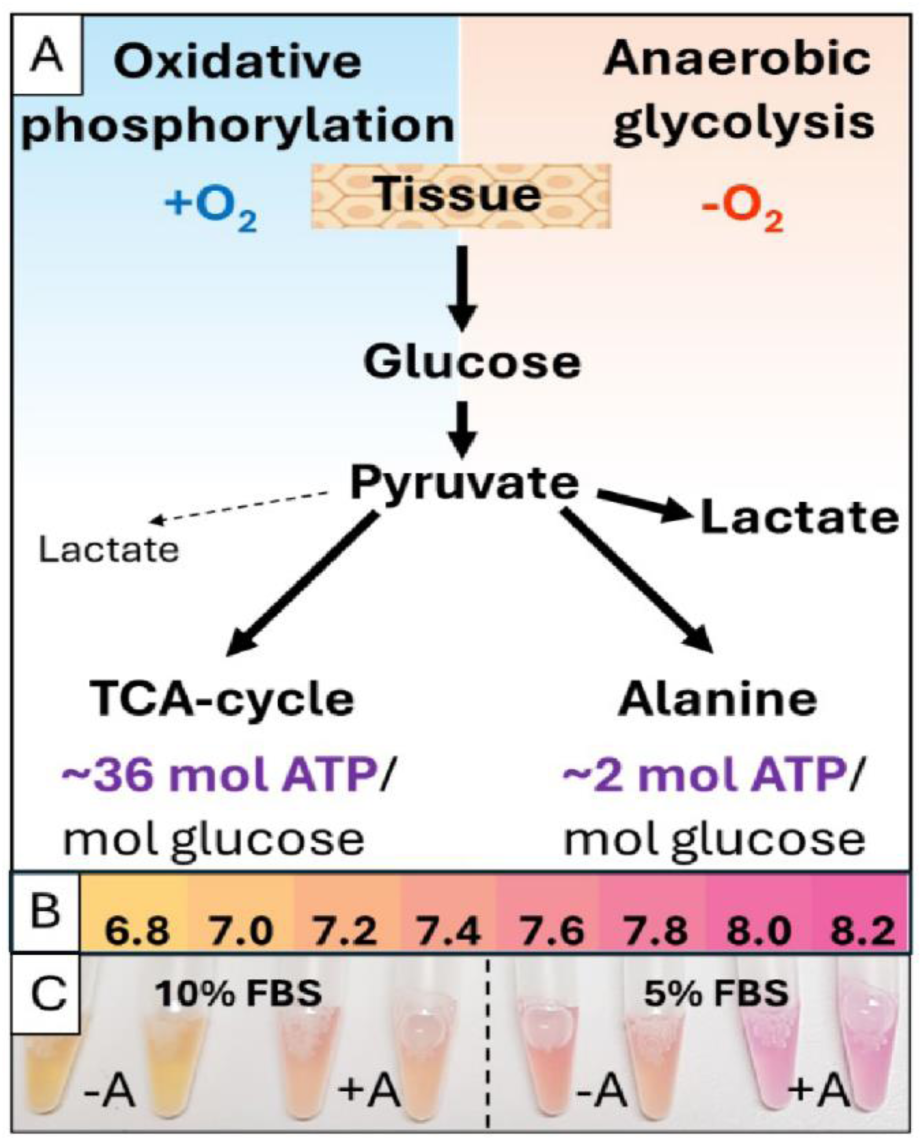
**(A)** Oxidative phosphorylation and anaerobic glycolysis as the major metabolic pathways to provide energy (Adenosine triphosphate (ATP)) to the cell and impact cell culture pH by lactate production. **(B)** Indicative colour chart (pH 6.8-8.2) of phenol red (Taylor Water Technologies LLC, K-1000 pH test). **(C)** C2C12 culture media colour at Day 6 with 10% or 5% FBS, co-cultured with (+A) or without (-A) *C. BDH-1* microalgae.

#### 2.7.2. Nuclear Magnetic Resonance (NMR) spectrum analysis of spent media

To obtain a more detailed understanding of the media composition when both cell types are co-cultured, we used NMR spectroscopy to investigate the chemical composition of the growth media after 6 days of growth with *C. BDH-1* or C2C12 cells, as well as C2C12-microalgae co-cultivation with 5% and 10% FBS (Fig. 5). Untargeted principal component analysis (PCA) of NMR spectra of the growth media from the four culture conditions after 6 days growth (Fig. 5A) showed clear systematic differences between mammalian cells only (blue) and algal co-cultures (green), as well as differences between cultures with 5% FBS (light colours) and 10% FBS (dark colours). The one-dimensional (1D) bivariate loadings plots of principal component 1 (PC1) and PC2 (Fig. 5B, C) allowed identification of the metabolites that are responsible for the sample differences observed in Fig 5A. Mammalian cell cultures exhibit higher levels of lactate, acetate, and alanine in the medium compared to the algal co-cultures (Fig 5B). Cultures with 5% FBS show higher levels of glucose, formate, phenylalanine, tyrosine, glutamate, glutamine, branched-chain amino acids, N-acetylated side chains, pyruvate, and succinate in the medium compared to cultures containing 10% FBS (Fig. 5C), which we attribute to the healthier growth in 10% FBS. We also observed ethylene glycol (EG) as a metabolite with altered levels in both PC1 and PC2 (Fig. 5B-C), likely being a contamination from one of the original media components.

**Figure 5:**
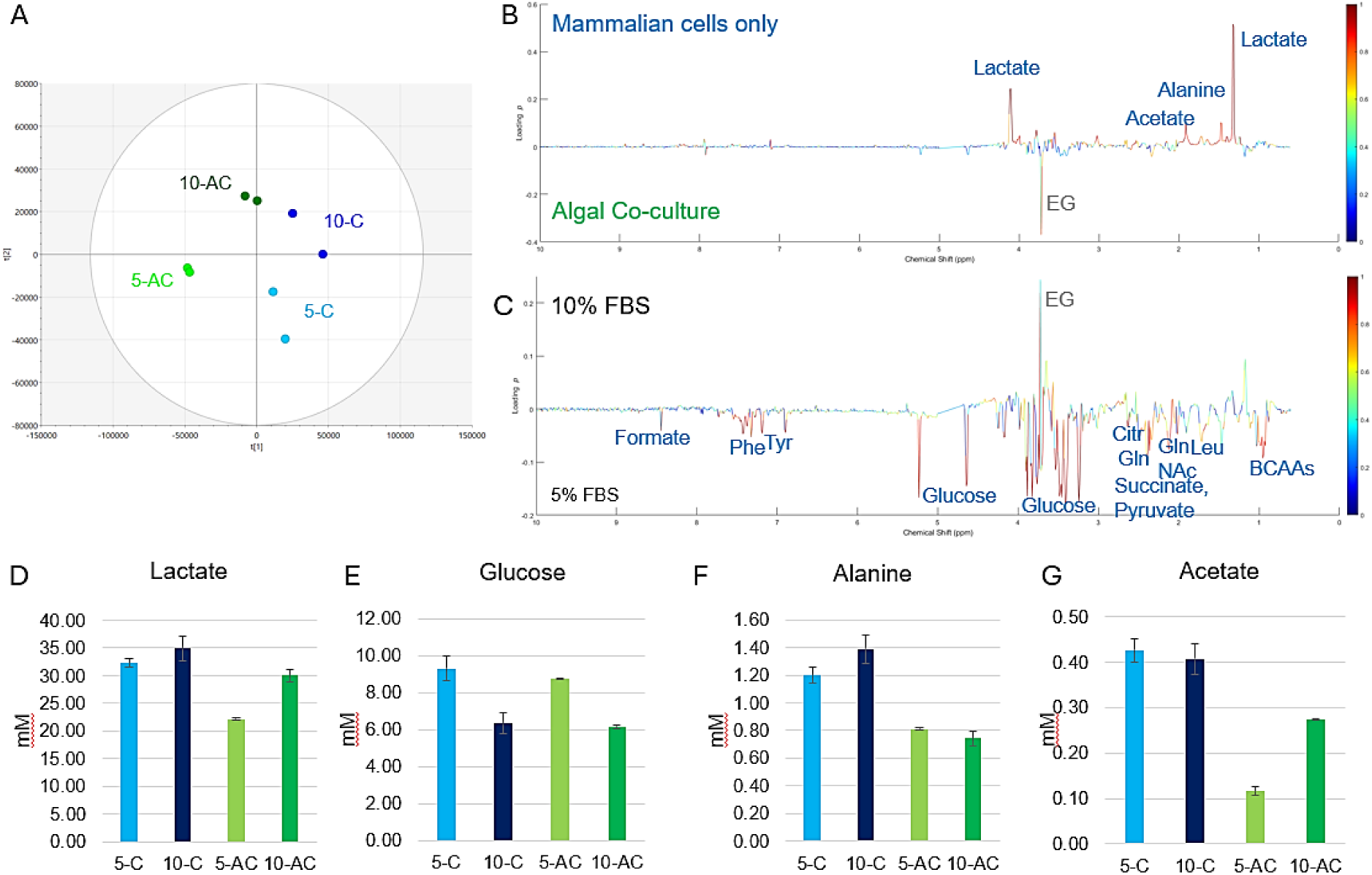
Multivariate statistical analysis with PCA of NMR spectra of mammalian cell culture and algal co-culture medium and targeted quantification of selected metabolites. **(A)** PCA Scores plot. Light blue: mammalian cell culture with 5% FBS (5-C), dark blue: mammalian cell culture with 10% FBS (10-C), light green: algal and mammalian cell co-culture with 5% FBS (5-AC), dark green: algal and mammalian cell co-culture with 10% FBS (10-AC). **(B)** 1D bivariate loadings plot of PC1, showing the differences between mammalian cell culture (top) and algal co-culture (bottom). The loadings coefficients for PC1 *p1* are shown as intensity of the line plot, and the absolute values of the correlation-scaled loadings coefficients |*p1(corr)|* as heatmap. **(C)** 1D bivariate loadings plot of PC2, showing the differences between 10% FBS (top) and 5% FBS (bottom). *p2* is shown as intensity and |*p2(corr)|* as heatmap. BCAAs: branched-chain amino acids, Citr: citrate, EG: ethylene glycol, Gln: glutamine, Leu: leucine, NAc: N-acetyl groups, Phe: phenylalanine, Tyr: tyrosine. **(D-G)** Quantification of the levels of lactate (D), glucose (E), alanine (F) and acetate (G) in the 4 culture conditions. The colour scheme and designation of culture conditions in panels (D-G) are the same as in panel (A).

Following this untargeted PCA, we quantified the NMR signals of the media components lactate, glucose, alanine and acetate in FBS and DMEM media after 6 days of growth with *BDH-1* or C2C12 cells, as well as C2C12-microalgae co-cultivation with 5 and 10% FBS (Fig. 5D-G, Table 3).

**Table 3:** Media acetate and glucose levels and metabolically active cell values at day 0 and 6 of cultivation in standard (10-C) and reduced FBS (5-C) conditions as well as co-cultivated with microalgae (10-AC, 5-AC).

| Sample |  | Lactate |  | Glucose |  | Average Cell Ratio |
| --- | --- | --- | --- | --- | --- | --- |
|  |  | [mM] | % | [mM] | % | % |
| Day 0 | DMEM pure | 0.8 | 2.3 | 25 | - |  |
|  | 10-DMEM start | 1.9 | 5.4 | 22.7 | **100.0** | - |
| Day 6 | 10-DMEM | 2.6 | 8.7 | 30.7 | 135.2 | - |
|  | 10-A | 2.2 | 6.3 | 27.5 | 121.1 | - |
|  | 10-C | 35 | **100.0** | 6.3 | 27.8 | **100** |
|  | 10-AC | 30 | 85.7 | 6.2 | 27.3 | 180 |
|  | 5-C | 32 | 91.4 | 9.3 | 41.0 | 30 |
|  | 5-AC | 22 | 62.9 | 8.7 | 38.3 | 55 |

##### 2.7.2.1. Lactate

We hypothesized that the maintained pink culture appearance is due to reduced levels of acidic fermentation products (especially lactate) in the culture medium when co-cultured with microalgae, which is in line with previous observations by Haraguchi et al.^(8)^. It is known that lactate suppresses oxidative phosphorylation^(33)^, has cytotoxic effects on animal cells and that lactate levels higher than 20 mM significantly decrease C2C12 cell viability^(34)^.

To confirm this hypothesis, we first quantified the lactate levels using known glucose concentrations as the reference integral. Interestingly, we already detected base levels of ∼2 mM lactate in the fresh unused DMEM media containing 10% FBS (10-DMEM) (Supplementary S2). To investigate the source of the lactate, pure DMEM as well as pure FBS (both as purchased) were analysed. The latter indeed showed significant amounts of lactate and thus was deemed the source of lactate in the freshly prepared 10-DMEM (Supplementary S2). Compared to fresh 10-DMEM, after 6 days we also detected an increase in the lactate levels in media without cells or in which only microalgae were cultivated (Sample 10-DMEM 6 days: 2.6 mM and Sample 10-A: 2.2 mM) (Table 3) which was attributed to component degradation. The analysis of the spent DMEM media from C2C12 cells that were co-cultivated with *BDH-1* cell (10-AC) showed increased lactate levels to ∼13.5-fold (∼30 mM) in comparison to the fresh (10-DMEM) or algae-only used DMEM media (10-A). However, as expected, the lactate levels in 10-AC were notably lower than in media from C2C12 cells alone (10-C, ∼35 mM) (Table. 3, Fig. 5D). A further decrease in lactate levels was observed in samples 5-AC, which showed lactate levels at ∼22 mM and thus just above levels that reportedly impact cell viability^(34)^.

Most microalgae do not possess genes to metabolize L-lactate and thus, similar to *C. BDH-1* in Fig. 2, cannot utilize L-lactate as a carbon source^(34)^. As microalgae cannot remove the lactate from the media (Fig. 2), we concluded that the algae caused reduced lactate production by the C2C12 cells. This is likely primarily due to the microalgal oxygen production allowing C2C12 cells to maintain oxidative phosphorylation with limited lactic acid production. Under oxygen-limited conditions, the cells switch to anaerobic glycolysis, and larger amounts of lactic acid are produced as a waste product^(31, 35)^ (Fig. 4A).

Remarkably, compared to 10-C samples on day 6, the samples 10-AC showed ∼80% higher numbers of metabolically active cells (Fig. 3B) at only 85% the lactate production (Table. 3). Overall, these findings confirm that reduced lactate levels may be responsible for the improved pH as visually indicated (Fig. 4C).

We thus consider that the standard frequent cell culture media changes are necessary rather due to harmful lactate waste accumulation than due to depletion of valuable nutrients, which are undesirably discarded in the process. This is supported by the observation, that microalgae co-cultivation doubles culture longevity and increases cell number, demonstrating that the media components necessary for cell growth are not the limiting factor (Fig. 3, Supplementary ST3).

##### 2.7.2.2. Glucose

To confirm that the microalgae do not compete for glucose with the C2C12 cells, we analysed the glucose levels in fresh media (10-DMEM) and Algae-only spent DMEM media (10-A4 and 10-A5) after 6 days, with both showing similar glucose levels (∼25 mM). This showed that the *BDH-1* did not compete with the C2C12 cells for the glucose necessary for their growth. In contrast to Chai et al.^(28)^ who reported *C. sorokiniana* UTEX 1230 to heavily utilize glucose up to a level of 88 mM, the data in Fig. 2C demonstrate that while it can utilize acetate, *BDH-1* metabolizes glucose poorly or not at all, as desired for our proposed application.

Notably, compared to fresh 10-DMEM or 6-day old 10-A4/10-A5 in which only microalgae were cultivated, media from co-cultivation (10-AC) as well as from C2C12 cells alone (10-C) both contained glucose levels of ∼25% (6 mM) (Table 3, Fig 5E). Glucose in 5-AC co-cultivation samples was only reduced to 35% (∼9 mM) of the standard 10-DMEM level.

Considering up to 80% more metabolically active C2C12 cells at both, days 2 and 6 in the co-cultivated samples 10-AC compared to the standard 10-C samples (Fig. 3A & D), these results indicate a more efficient glucose utilization through co-cultivation as expected from energy production using oxidative phosphorylation (Supplementary ST3).

##### 2.7.2.3. Alanine

In line with the hypothesis that these microalgae support oxidative phosphorylation in the C2C12 cells, we have also noted that media from co-cultivation contained less alanine, with 5-AC samples containing the least (Fig. 5F). During anaerobic glycolysis, muscle cells degrade amino acids for energy needs. The resulting nitrogen is then transaminated to pyruvate forming alanine, which is then exported from the cell^(36)^. Under oxidative phosphorylation, cells typically do not excrete significant amounts of alanine as pyruvate is funnelled into the TCA cycle rather than converted to alanine (Fig. 4A). It has recently been proposed that alanine supplementation of the microalgae *Chlorella pyrenoidosa* increases biomass yields and lipid production^(37)^; it is therefore possible that the here introduced microalgae *C. BDH-1* is also able to use alanine. However, given the reduced lactate concentrations in the co-cultures and the microalgal inability to metabolize lactate, it is more likely that the C2C12 cells in the presence of microalgae produce less alanine due to photosynthetic oxygen provision.

##### 2.7.2.4. Acetate

We have identified 1.5-fold higher acetate levels in DMEM media from C2C12 cell cultivation (10-C) compared to levels in co-cultivation media with 10% FBS after 6 days (10-AC). The lowest acetate levels were found in co-cultivation media with only 5% FBS (5-AC, Fig. 5G). Acetate has been shown to negatively impact cell viability in cancer cells *in vitro*^(38)^, and its production from glucose and excretion into the media has been coupled to mitochondrial metabolism and a phenomenon that becomes more pronounced with nutritional excess (e.g. hyperactive glucose metabolism)^(39)^. As we have shown that *BDH-1* prefers acetate as a carbon source (Fig. 2D), the microalgae likely consume acetate from the media, which aids in the microalgal-positive impact on the media longevity. It is also likely that microalgal oxygen supply supports mitochondrial metabolism and thus reduces acetate secretion. While the specific mechanism by which the *C. BDH-1* reduces acetate levels has not been conclusively determined, the reduced acetate levels themselves confirm our findings that microalgae improve the culture pH (Fig. 4C) and reduce waste products in the media, improving cell culture longevity.

### 2.8. *Chlorella BDH-1* as a candidate for medical tissue culture applications

While industry-scale applications such as cultivated meat address a larger market, laboratory-scale 3D tissue cultivation is importance in the medical sector for regenerative therapy applications such as custom cell-sheet cultivations for transplants^(2)^, or future-oriented organoid transplants^(3)^, and photoautotrophic tissue engineering^(40)^, all seeking animal-free and more accessible applications. They therefore provide a more imminent application platform for the development of tissue co-cultivation with microalgae. To determine whether 3D tissue co-cultivation with *BDH-1* may be safe for future patient applications, we assessed live algal immunogenicity using primary BMDM. First, we assessed the cell viability of the macrophages at 4 h and 24 h post-treatment with different ratios of BMDM to microalgae in pure media, as well as with the inflammatory stimulus lipopolysaccharide (LPS) as a positive control. The microalgae did not show any adverse effects. All samples with microalgae potentially show viability improvement at 24 h compared to samples without microalgae in pure media as well as LPS controls (Fig. 6A, Supplementary ST4). Next, we measured secreted levels of the cytokine Interleukin 6 (IL-6) (Fig.6B, Supplementary ST5), a proinflammatory cytokine used as an inflammation marker. It was confirmed that after 24 h, *BDH-1* did not induce an inflammatory response as assessed by IL-6 production, which the literature supports^(41-43)^. Consequently, no adverse effects of microalgae (e.g. when co-cultured with skin grafts) are expected, although a more comprehensive assessment will be required.

**Figure 6:**
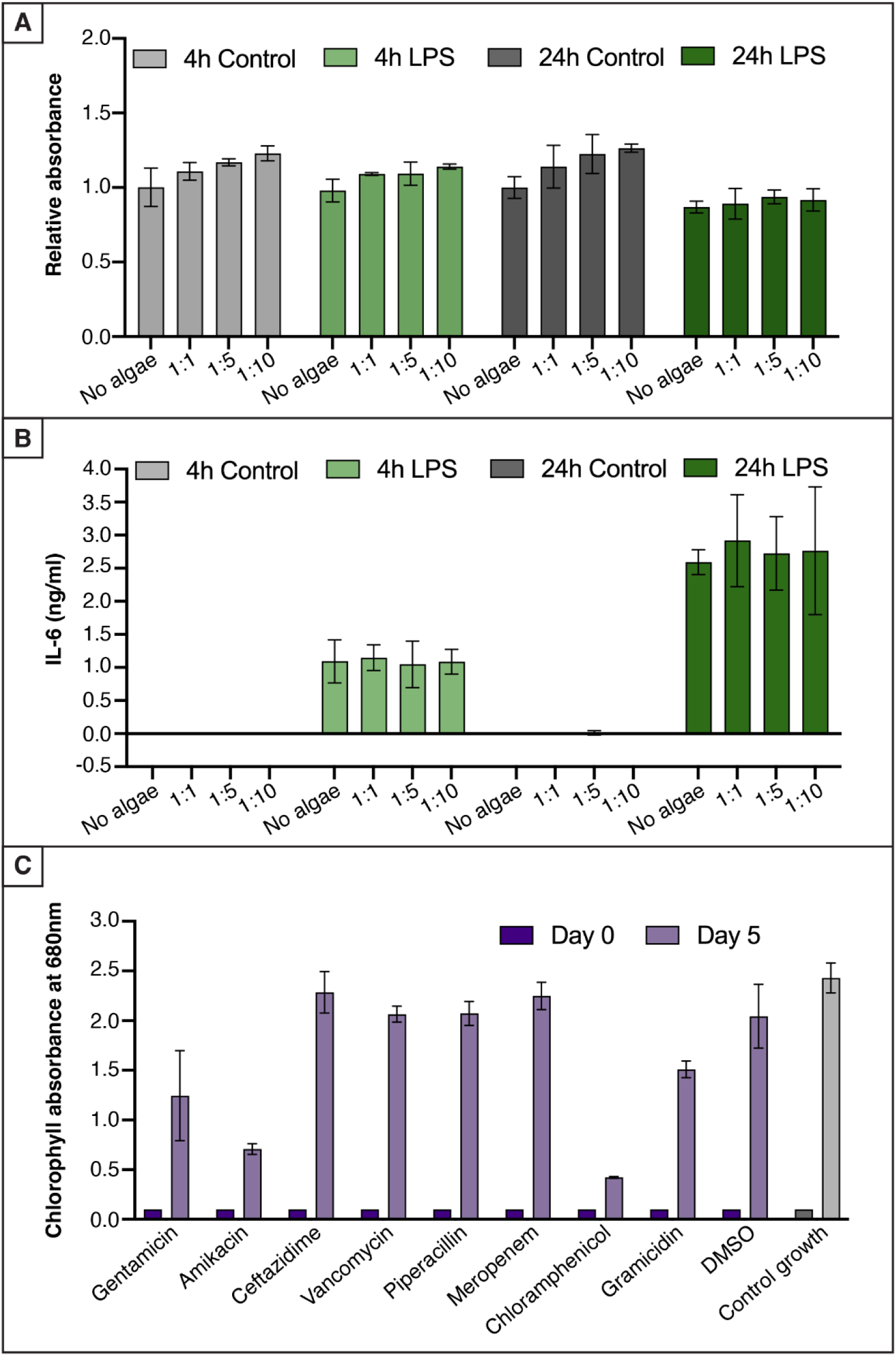
*Chlorella BDH-1* suitability tests for medical applications. **(A)** Relative MTT cell viability assay of mouse BMDMs treated ± different ratios of microalgae (1:1, 1:5 and 1:10) in media alone or with the lipopolysaccharide (LPS) as positive control. **(B)** Secreted levels of the inflammatory cytokine interleukin-6 (IL-6) in response to different ratios of BMM to microalgae. **(A) and (B):** Quantitation is relative to “no algae” control (mean±SEM) from two independent experiments with two technical replicates each. **(C)** *C. BDH-1* growth (mean±SD, n=2) after 5 days of exposure to clinically relevant antibiotics at 1× maximum patient serum concentration as derived from MIMS database.

The absence of inflammatory response triggered by *BDH-1* makes it an ideal candidate for photoautotrophic tissue development and applications in patients. We therefore tested its susceptibility to 6 different classes of clinically relevant antibiotics (Table 4) at concentrations between 1× to 8× peak patient serum concentration (Fig. 6D, Supplementary S3) as derived from MIMS (Monthly Index of Medical Specialties) pharmaceutical database (https://www.mims.com.au/).

**Table 4:**
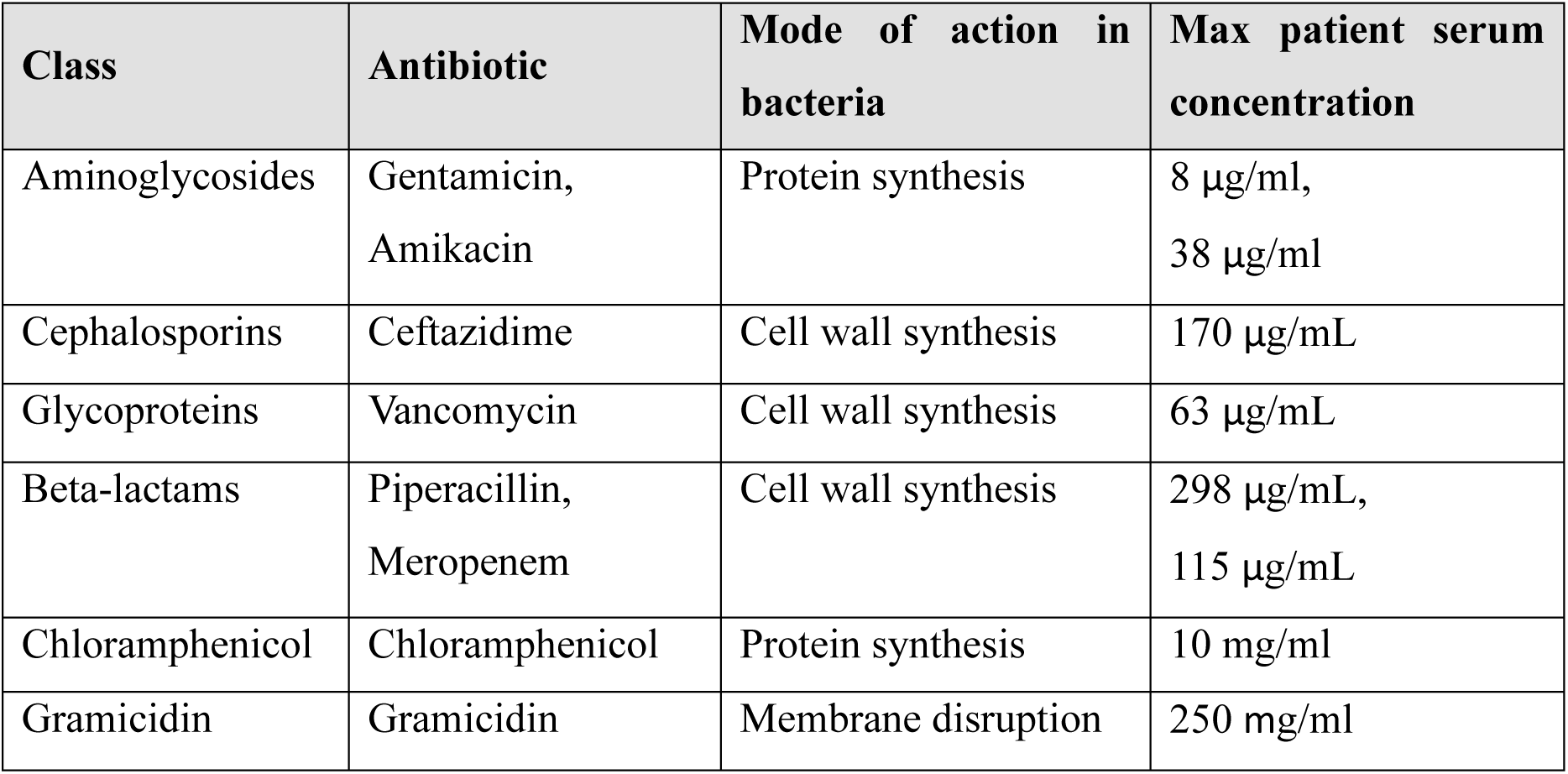
Antibiotics tested in this study, their mode of action and maximum peak serum concentrations as per MIMS.

| Class | Antibiotic | Mode of action in bacteria | Max patient serum concentration |
| --- | --- | --- | --- |
| Aminoglycosides | Gentamicin,<br>Amikacin | Protein synthesis | 8 µg/ml,<br>38 µg/ml |
| Cephalosporins | Ceftazidime | Cell wall synthesis | 170 µg/mL |
| Glycoproteins | Vancomycin | Cell wall synthesis | 63 µg/mL |
| Beta-lactams | Piperacillin,<br>Meropenem | Cell wall synthesis | 298 µg/mL,<br>115 µg/mL |
| Chloramphenicol | Chloramphenicol | Protein synthesis | 10 mg/ml |
| Gramicidin | Gramicidin | Membrane disruption | 250 mg/ml |

*BDH-1* growth was observed for 5 days and recorded as chlorophyll absorbance at 680 nm. Green microalgal chloroplasts, although lacking peptidoglycan, are evolutionarily derived from cyanobacteria^(44)^ and are therefore susceptible to antibiotics targeting the prokaryotic cell machinery. As expected, antibiotics that interfere with prokaryotic protein synthesis (Gentamicin, Amikacin and Chloramphenicol) inhibited algal growth at 1× peak patient serum concentration (Fig. 6D) with a dose-dependent increase in severity up to 8× peak concentrations (Supplementary S3). Antibiotics that addressed cell wall synthesis and bacterial-membrane integrity did show little to no effect on microalgal growth (Fig. 6D).

The chosen antibiotic concentrations are between 1× - 8× peak concentrations in patients’ serum and therefore unlikely to directly affect *BDH-1* for any external application. However, the demonstration of *BDH-1* being unaffected by a large number of clinically relevant antibiotics opens up the possibility of direct medical application of *C. BDH-1* e.g. for chronic wound oxygenation^(45)^.

## 3. CONCLUSION

The successful co-cultivation of microalgae with mammalian cells for laboratory, industrial or medical applications requires the careful selection of a microalga that can provide a solid support system within the more sensitive mammalian cell culture conditions. We have screened several different microalgae species to identify a microalgal candidate that is suitable for mammalian cell growth conditions in terms of temperature and salinity while not competing with the mammalian cells for glucose as an energy source. We identified *C. BDH-1,* a locally isolated Australian microalgae, to perform best in all tested temperature and salinity conditions. We also demonstrated that *C. BDH-1*, unlike other reported *Chlorella* strains, does not utilize glucose for growth, showing a similar profile to photoautotrophic cultivation under glucose provision and thus not competing with the mammalian cells for glucose. Considering potential future medical applications, we showed that *C. BDH-1* does not elicit an inflammatory response as assessed by IL-6 production and is not susceptible to several clinically relevant antibiotics. We therefore conclude that *C. BDH-1* provides an ideal candidate for mammalian cell-microalgae co-cultivation.

C2C12 are metabolically demanding and temperature-sensitive mouse myoblast cells. Here, we established that C2C12 culture longevity and productivity improve by successful co-cultivation with *C. BDH-1* cells at an optimized ratio for both cell types (Fig.3), similar to those generally observed in symbiotic relationships^(46)^. Visual pH assessment (Fig. 4) and NMR spectroscopy of the media components indicated that key reasons for improved longevity and boosted cell viability were more efficient glucose utilization and thus reduced lactate and acetate production leading to reduced lactate and acetate accumulation in the media (Fig. 5), likely in parallel to microalgal acetate consumption. It is plausible that by providing photosynthetic oxygen the microalgae facilitate maintenance of oxidative phosphorylation allowing more effective glucose usage and reducing the accumulation of waste products. This improved longevity, as well as higher viable cell numbers, and indicates that frequent media changes in culture systems (which lack a cardiovascular system as an oxygen supply and for waste removal), are required due to waste product accumulation rather than exhaustion of growth resources. Co-cultivation therefore has the potential to reduce media costs by allowing the cells to utilize the remaining growth resources which are usually discarded.

Co-cultivation of C2C12 cells with microalgae also allowed a 50% reduction in costly and ethically concerning FBS while, to an extent, maintaining improved longevity in comparison to standard cultivation procedures in DMEM with 10% FBS, which itself introduces the culture waste product, lactate.

In summary, we have shown that co-cultivation of mammalian cells with our *Chlorella BDH-1* is beneficial for mammalian cells and thus has the potential to reduce media costs for laboratory and larger-scale cell culture systems and to provide a suitable and safe candidate to explore medical applications.

## 4. MATERIALS & METHODS

### 4.1. Cell lines, culture media, and cell maintenance

#### 4.1.1. Microalgae cells

Microalgae strains were obtained from the Chlamydomonas Resource Center in Minnesota, USA (*Chlamydomonas reinhardtii* CC124, CC125, CC1883), CSIRO Australian National Algae Supply Service (ANASS) (*Chlorella vulgaris*, *Nanochloropsis oceanica* (Droop) Green (CS-179), *Dunaliella salina* CS-265, *Phaeodactylum tricornutum* CS-29), while *Chlorella sp. 11H5*^(19)^, and *Chlorella BDH-1* were locally isolated.

For maintenance, microalgae were cultivated either in liquid TAP medium^(23)^ for freshwater strains, or F/2 medium^(25)^ for saltwater strains under continuous warm white LED-light (150 µmol photons m^-2^ s^-1^), atmospheric CO_2_ either at room temperature (20 °C) or at 37 °C, as indicated.

#### 4.1.2. Mammalian cells

C2C12 cells were obtained from the American type culture collection (ATCC CRL-1772) and routinely grown in Dulbecco’s Modified Eagle Medium (DMEM, Gibco™ DMEM 11995065) supplemented with 10% foetal bovine serum (FBS – French Origin, Bovogen Biologicals), 2 mM L-Glutamine (Gibco™) and 100 µg/mL Ampicillin (Novachem) at 37°C and 5% CO_2_. Concentrations of FBS were varied in percentage as indicated in each experiment.

C57Bl/6 mice aged between 8 to 12 weeks were used to generate primary mouse bone marrow-derived macrophages (BMDM). Murine bone marrow cells were extracted from femurs and tibias of mice and cultured for 7 days in RPM1 media (Gibco, Thermofisher), containing Glutamax (2 mM, Life Technologies, Waltham, Massachusetts, USA), 10% FCS (Gibco, Thermofisher, USA), 20 U/mL penicillin, 20 g/mL streptomycin (Gibco, Thermofisher) and 150 ng/mL recombinant human CSF-1 (generated by The University of Queensland Protein Expression Facility).

### 4.2. Algae growth assessment in different media

For microalgal growth assessments different media, PCM^(24)^, F/2^(25)^ and TP media^(23)^) supplemented with acetate (18 mM, 36 mM), glucose (25 mM, 50 mM) and lactate (18 mM, 36 mM), as well as DMEM (Gibco™ DMEM 11995065) with 10% FBS (FBS – French Origin, Bovogen Biologicals) were used. Algae were inoculated and grown at room temperature and 37 °C with 150 µmol photons m^-2^ s^-1^ continuous warm white LED-light. Algae growth was measured based on chlorophyll absorption (680 nm) with a spectrophotometric plate reader (Tecan Infinite® M Plex).

### 4.3. Sequencing

To identify species and strains of isolated *BDH-1*, 18S and 16S sequencing was performed. The microalgae were cultivated in TAP media to log-phase. Cells were harvested, resuspended in DNA extraction buffer [1 M KCl, 0.01 M EDTA, 0.1 M Tris-HCl, pH 9.5], and disrupted at 12 ksi using a Continuous Flow Cell Disruptor CF1 (Constant Systems). Genomic DNA was extracted according to Thomson and Henry^(47)^. Two separate sections of the 18S ribosomal RNA gene, together spanning a large portion of the gene, were sequenced using primer pairs: 18S-F1: 5’ CCTGTCTCAGATTAGCC 3’ and 18S-R1: 5’ ACCAGACTTGCCCTCC 3’, 18S-F2: 5’ GCGGTAATTCCAGCTCCAATAGC 3’ and 18S-R2: 5’ GACCATACTCCCCCCGGAACC 3’. Subsequently *Chlorella* specific 16S primers 16S-F: 5’ GGGATTAGATACCCCCTGTAGTCCT 3’ and 16S-R: 5’ AGGACTACAGGGGTATCTAATCCC 3’ in combination with universal primers^(30)^ were used to sequence the chloroplast RNA gene. Obtained results were assessed for similarities to known sequences using NCBI Basic Local Alignment Search Tool (BLAST)^(29)^.

### 4.4. Co-cultivation of C2C12 and microalgae cells

For co-cultivation experiments, Corning® Transwell® 24-well plates with polycarbonate membranes were used to separate microalgae from mammalian cells. Experiments were performed in a standard tissue culture incubator at 37 °C, 5% CO_2_. A commercially available warm white LED strip was used to provide continuous 100 µmol photons m^-2^ s^-1^ illumination. 20,000 C2C12 cells were seeded into each well in a total of 500 µl of DMEM growth media (containing 5% or 10% FBS as indicated). Cells were allowed to settle for 24 h. Microalgae cells were resuspended in DMEM growth media with the FBS concentrations matching those used for the C2C12 cells, and 150 µl of microalgae cell suspension was added to transwells at mammalian:microalgae cell ratios of 1:200, 1:20, 1:2, 1:0.2 and 1:0.02. Mammalian cell growth was monitored visually and by assessing cell viability with resazurin assays at days 2 and 6.

### 4.5. Cell viability assays

For the resazurin assays, a 100 mM resazurin solution was prepared in PBS. Resazurin is a soluble dye that is reduced to highly fluorescent resorufin in proportion to the metabolic activity of a cell population^(48)^. After removal of the algae-containing transwell, resazurin solution was added to the 500 µl DMEM media to 10 mM final concentration. After 3 h of incubation at 37 °C in the dark, resazurin fluorescence was measured with a plate reader (Tecan Infinite® M Plex) at 590 nm using 560 nm excitation. PrestoBlue™ Cell Viability Reagent (Thermo Fisher Scientific) was also used with a 1 h incubation time, 545 nm excitation, and 600 nm emission. For MTT (3-[4,5-dimethylthiazol-2-yl]-2,5 diphenyl tetrazolium bromide) assays, BMM cultured with or without microalgae were incubated with 1 mg mL^−1^ MTT reagent (Sigma-Aldrich), diluted in complete BMM media without antibiotics. Cells were left at 37 °C for 1–3 h. The MTT media was removed, and formazan crystals were dissolved in 300 μl 100% isopropanol. Once the formazan precipitate was fully dissolved, the dissolved formazan was transferred to a 96-well plate and the absorbance at 510 nm was read in an Infinite M Plex (Tecan, Mannedorf, Switzerland) plate reader.

### 4.6. Growth media component analysis using nuclear magnetic resonance (NMR) spectroscopy

To analyse the effect of microalgae on mammalian cell growth, cell culture supernatant from C2C12 cells was assessed after 6 days of cultivation in DMEM growth media (containing 5% or 10% FBS), both with and without microalgae co-cultivation. Additional controls included: unsupplemented DMEM media, FBS alone, and DMEM growth media with 5% or 10% FBS, respectively. NMR samples were prepared by mixing 160 μL medium with 20 μL 1.5 M potassium phosphate buffer pH 7.4 and 20 μL of a solution containing 1 mM sodium 2,2-dimethyl-2-silapentane-5-sulfonate-d_6_ (DSS) as a chemical shift reference, and 1 mM difluorotrimethylsilanylphosphonic acid (DFTMP) as an internal pH indicator in D_2_O, yielding a final sample volume of 200 μL with final concentrations of 100 μM DSS, 100 μM DFTMP, and 7.5% D_2_O. Samples were transferred into 3-mm NMR tubes for measurement.^1^H NMR spectra were recorded on a Bruker 800 MHz Avance IIIHD NMR spectrometer (Bruker Biospin, Rheinstetten, Germany) operating at a ^1^H frequency of 800.26 MHz and equipped with a 5 mm self-shielded z-gradient triple resonance probe and a chilled SampleJet sample changer. For each sample a 1D Carr-Purcell-Meiboom-Gill (CPMG) spectrum was acquired at 298 K with the *cpmgpr1d* pulse sequence [RD-90°-(τ-180°-τ)*_n_*-acq] (Bruker Biospin pulse program library). The transmitter frequency was set to the frequency of the water signal, and water suppression was achieved by continuous wave irradiation during the relaxation delay (RD) of 4.0 s. A fixed spin−spin relaxation delay 2nτ of 75.6 ms (τ = 300 μs) was used to eliminate the broad signals from high molecular weight analytes. After 4 dummy scans, 128 transients were collected into 73,728 data points using a spectral width of 20 ppm, leading to a total experiment time of 18.9 min per spectrum. All spectra were processed using TOPSPIN version 4.3.0 (Bruker Biospin, Rheinstetten, Germany). The free induction decays (FIDs) were multiplied by an exponential window function with a line broadening factor of 0.3 Hz before Fourier transformation, manual phase and baseline correction. The resulting spectra were referenced to the glucose anomeric doublet at δ = 5.233 ppm. The ^1^H NMR spectra were data-reduced in with an in-house script in Matlab (MathWorks, Natick, MA, USA) to consecutive integral regions of 0.01 ppm width (“buckets”), covering the range of δ = 10.0–0.06 ppm. The chemical shift region at δ = 5.0–4.7 ppm was excluded to eliminate the effects of imperfect water suppression. Subsequently, the bucketed data matrices were imported into the SIMCA 18 software package (Sartorius Stedim AB, Umeå, Sweden) for multivariate statistical analysis. The data matrix with the bucketed 1D spectra (*n*=8 samples, *k*=910 variables) was Pareto scaled and analysed with principal components analysis (PCA) to investigate inherent differences in the samples^(49)^. In SIMCA, the number of latent components (*A*) for the PCA model was optimized by cross validation. *R*^2^*X* and *Q*^2^ were used to evaluate model quality. *R*^2^*X* is the fraction of the sum of squares explained by the latent components in the model, representing the variance of the *X* variables, and *Q*^2^ is the predictive ability parameter of the model, which is estimated by cross validation. The figures of merit of the PCA model are *A*=3, *R^2^X*=0.959, *Q^2^*=0.849. A scores plot was used to interpret the PCA model, together with bivariate 1D loadings plots, in which loadings coefficients *p* were plotted against the chemical shift values of their respective variables, and the correlation-scaled loadings coefficients |*p*(*corr*)| were superimposed on the loadings plot as a heatmap color scale^(50)^. These plots contain the same information as a traditional S-plot, which indicates which variables influence the model with high reliability and are of relevance in the search for significantly altered metabolites.

The metabolites observed to change significantly in the multivariate analysis were identified with Chenomx NMR Suite 7.0 (Chenomx Inc., Edmonton, Canada), and selected metabolites were then quantified by peak integration of the spectral region(s) containing non-overlapped signals of the respective metabolite using the AMIX integration tool on the full-resolution spectra. Only metabolite signals with minimal peak overlap were used in this analysis.

### 4.7. Inflammatory response assessment

#### 4.7.1. Macrophage – Algae co-culture

On day 6 of differentiation, BMDM were harvested and plated at a density of 0.2×10^6^ cells per well in complete RPMI media without antibiotics and left to adhere overnight. Cells were cultured with algae at different ratios as noted in the figures as macrophage:algae cell ratios. The TLR4 agonist LPS, chromatographically purified from *Salmonella enterica* serotype Minnesota Re 595 (Cat: L2137, Sigma-Aldrich) was used at the concentrations listed in individual figures. Cell supernatants were collected at 4 h and 24 h post-treatment to be analyzed for inflammatory cytokine expression via ELISA. The remaining attached cells were then used for MTT assays.

#### 4.7.2. Enzyme-linked immunosorbent assay (ELISA)

Rat anti-mouse IL-6 (CAT:554400) and Rat biotinylated anti-mouse IL-6 (CAT:554402) antibodies were used to assess levels of secreted mouse IL-6 via sandwich ELISA. All antibodies were obtained from BD Bioscience, California, USA. Briefly, 96-well ELISA plates (Nunc, Rochester, NY, USA) were coated with capture antibody (diluted in 0.1 M sodium bicarbonate, pH 8.35) overnight. Plates were washed with PBS containing 0.05% Tween before being blocked with 10% FBS in PBS for 2 h at 37°C. Plates were washed before samples and standards (diluted in the BMM complete media) were added and incubated for 2 h at 37°C. Plates were then sequentially incubated and washed with secondary antibody (diluted in 10% FBS in PBS) for 1 h at 37°C, followed by extra-avidin (1:1000 dilution in 10% FBS in PBS) for 20 min at 37°C. After further washing, the TMB (3,3’,5,5’-tetramethylbenzidine) substrate (BD OptEIA; BD Biosciences) was added. Reactions were stopped using 2 M sulfuric acid and absorbance at 450 nm was read using a plate reader (Infinite M Plex, Tecan). Cytokine levels were calculated by extrapolation from a sigmoidal curve analysis of the standards.

### 4.8. Antibiotic compatibility assessment

The effect of antibiotics on microalgae *C. BDH-1* growth was evaluated by incubation for 96 h at 37 °C and 150 µmol photons m^-2^ s^-1^ in 96-well PS plates (Corning 3370). Clinically relevant antibiotics were selected, and concentrations at 8×, 4×, 2× and 1× peak serum concentration in patients (Monthly Index of Medical Specialties) were tested in duplicate. For chloramphenicol and gramicidin, which are typically applied topically, concentrations were based on the concentrations in typical ointments. The following antibiotics were tested at these final concentrations: gentamicin at 8, 16, 32, 64 µg/mL, amikacin at 37.5, 75, 150, 300 µg/mL, ceftazidime at 168.75, 337.5, 675, 1350 µg/mL, vancomycin at 62.5, 125, 250, 500 µg/mL, piperacillin at 300, 600, 1200, 2400 µg/mL, meropenem at 112.5, 225, 450, 900 µg/mL, chloramphenicol at 1250, 2500, 5000, 10000 µg/mL and gramicidin at 250, 500, 1000, 2000 µg/mL.

### 4.9. Statistical data analysis of C2C12 co-cultivation experiments

To enable comparability across all biological replicates (independent experiments), obtained data was standardized by setting each experimental *no algae* control=1. For each biological replicate, the result was estimated as the mean value of the technical replicates. For each biological replicate, the variance – and therefore the standard deviation - was calculated to indicate the degree of dispersion, or error. The final overall result was then estimated as the mean across all biological replicates. The overall standard deviation across all biological replicates was also computed to indicate the error of the final estimate. This was achieved by averaging the variances of all the individual replicates. Number and values of each independent biological replicate as well as the overall results and standard deviation are indicated in Supplementary Tables ST1 and ST2.

## SUPPLEMENTARY DATA

**Table ST1:**
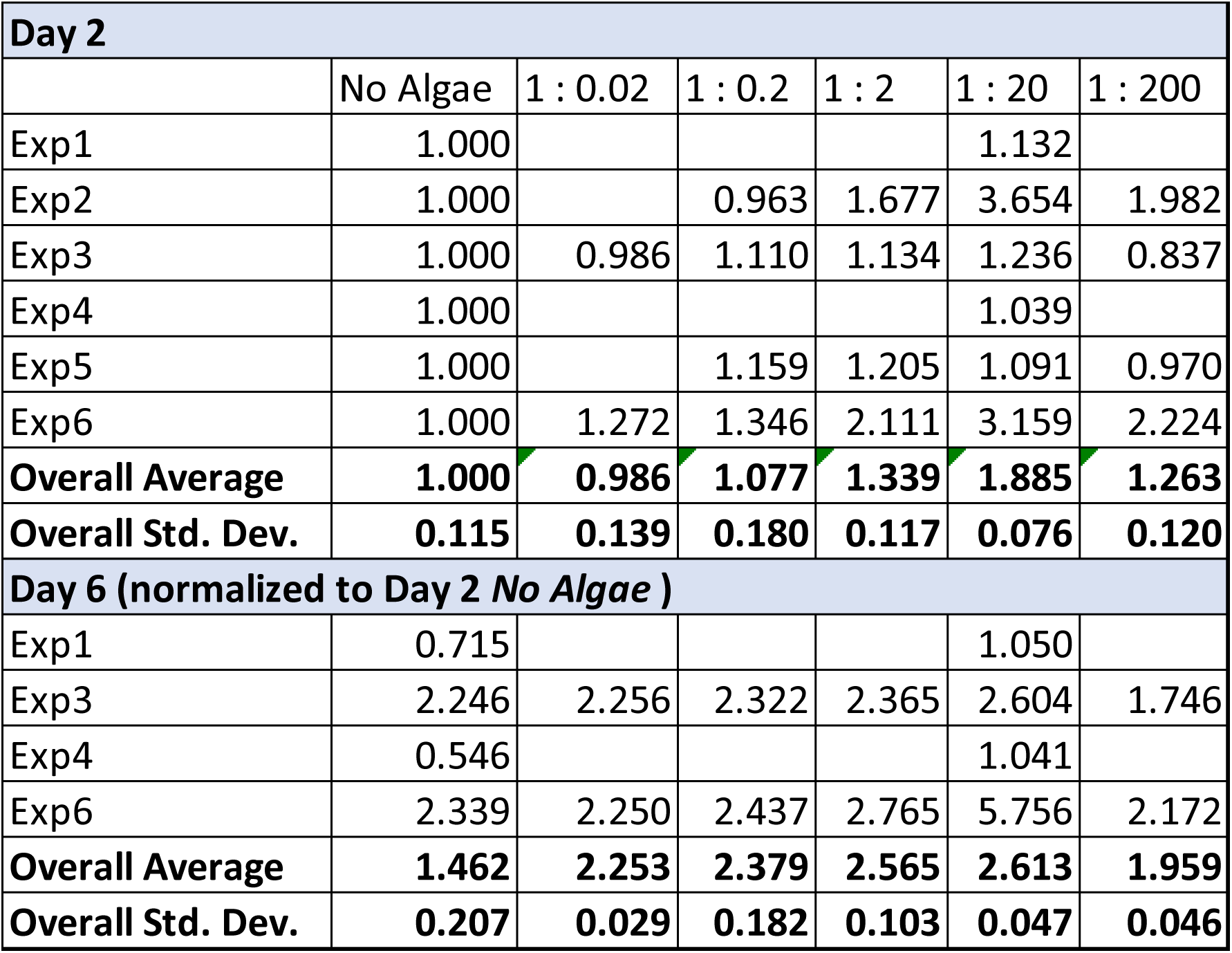
Independent experiments (Exp) for resazurine fluorescence data from varying mammalian : algae cell co-cultivation ratios (mean±SD, n=3).

**Table ST2:**
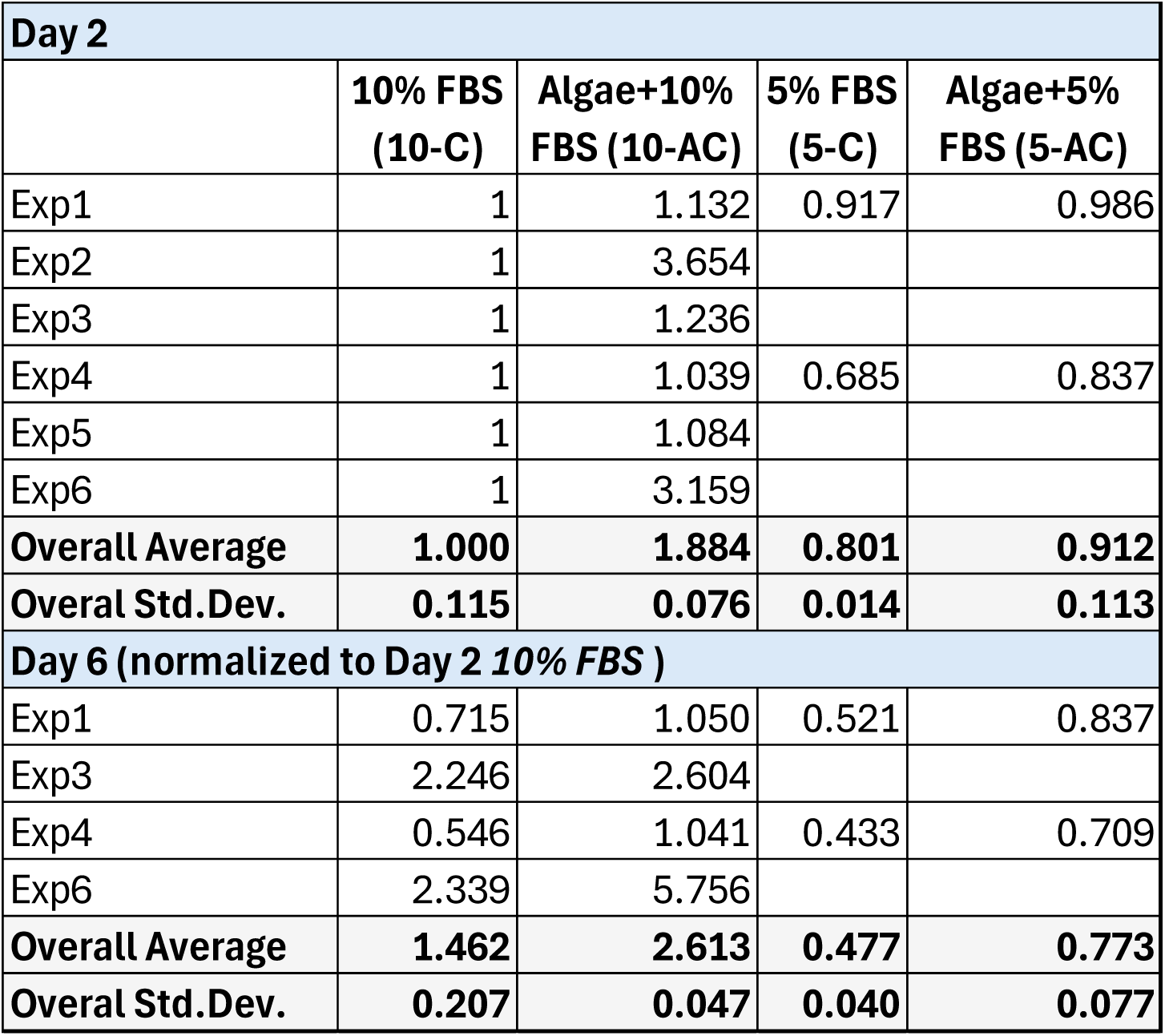
Independent experiments (Exp) to obtain relative fluorescence data normalized to control (10% FBS, 10-C) at day 2 from C2C12 cultivation with or without microalgae in 10% or 5% FBS, respectively (mean±SD, n=3).

**Figure S1:**
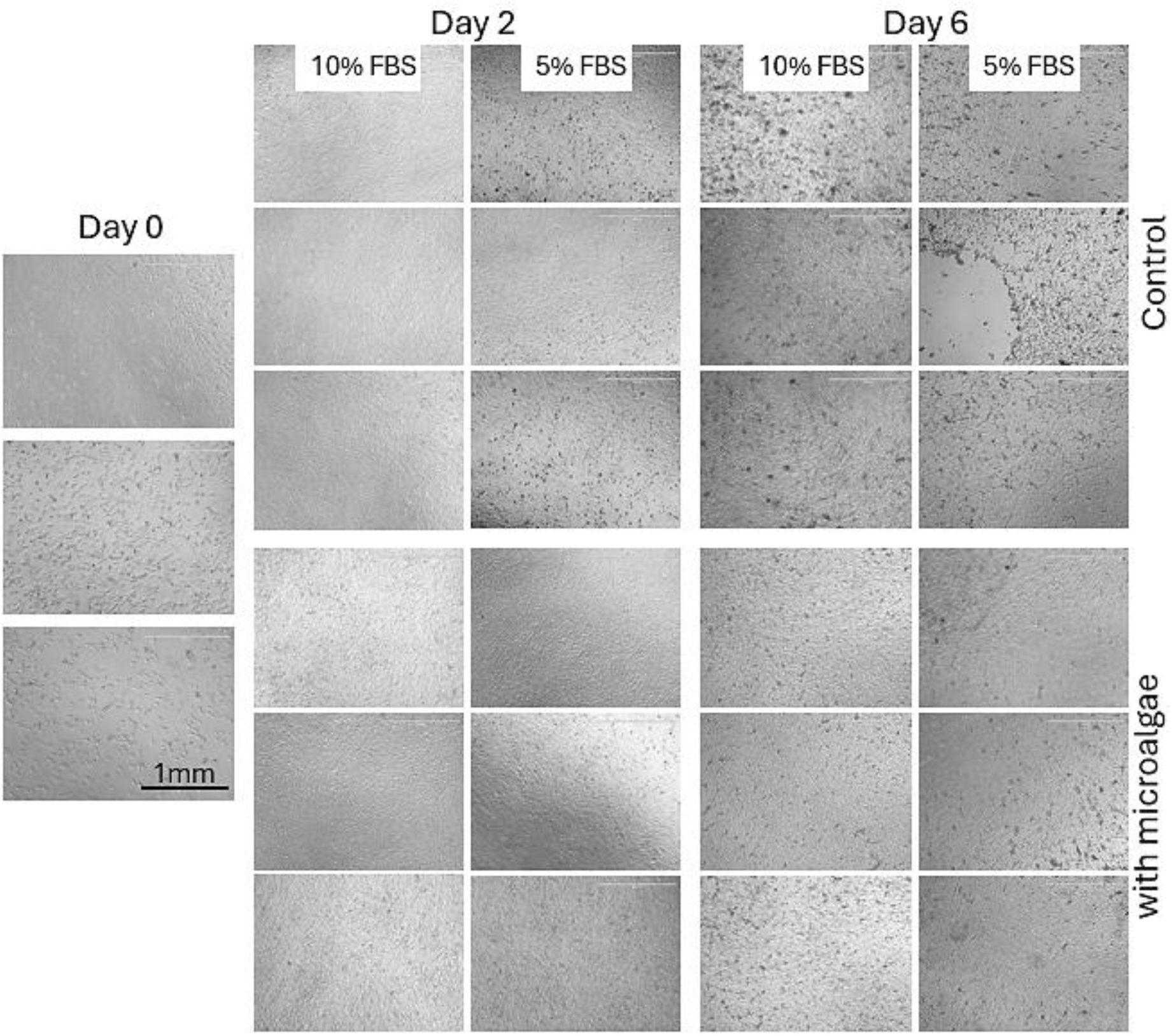
Brightfield images of C2C12 cells at 4× magnification on day 0, day 2, and day 6 cultivated with 5% or 10% FBS with or without microalgae.

**Figure S2:**
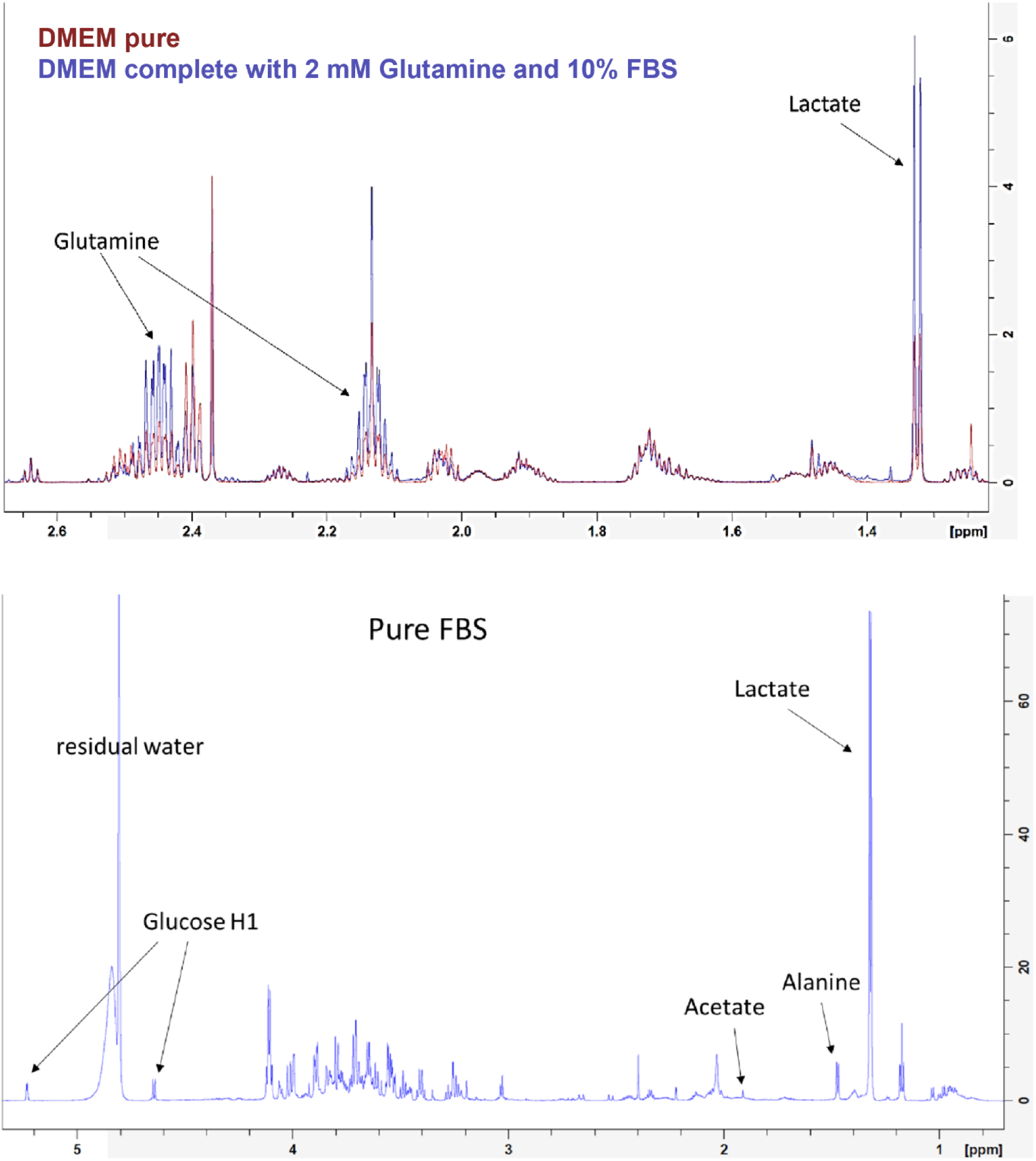
NMR spectra of DMEM pure and complete, as well as pure FBS.

**Table ST2:**
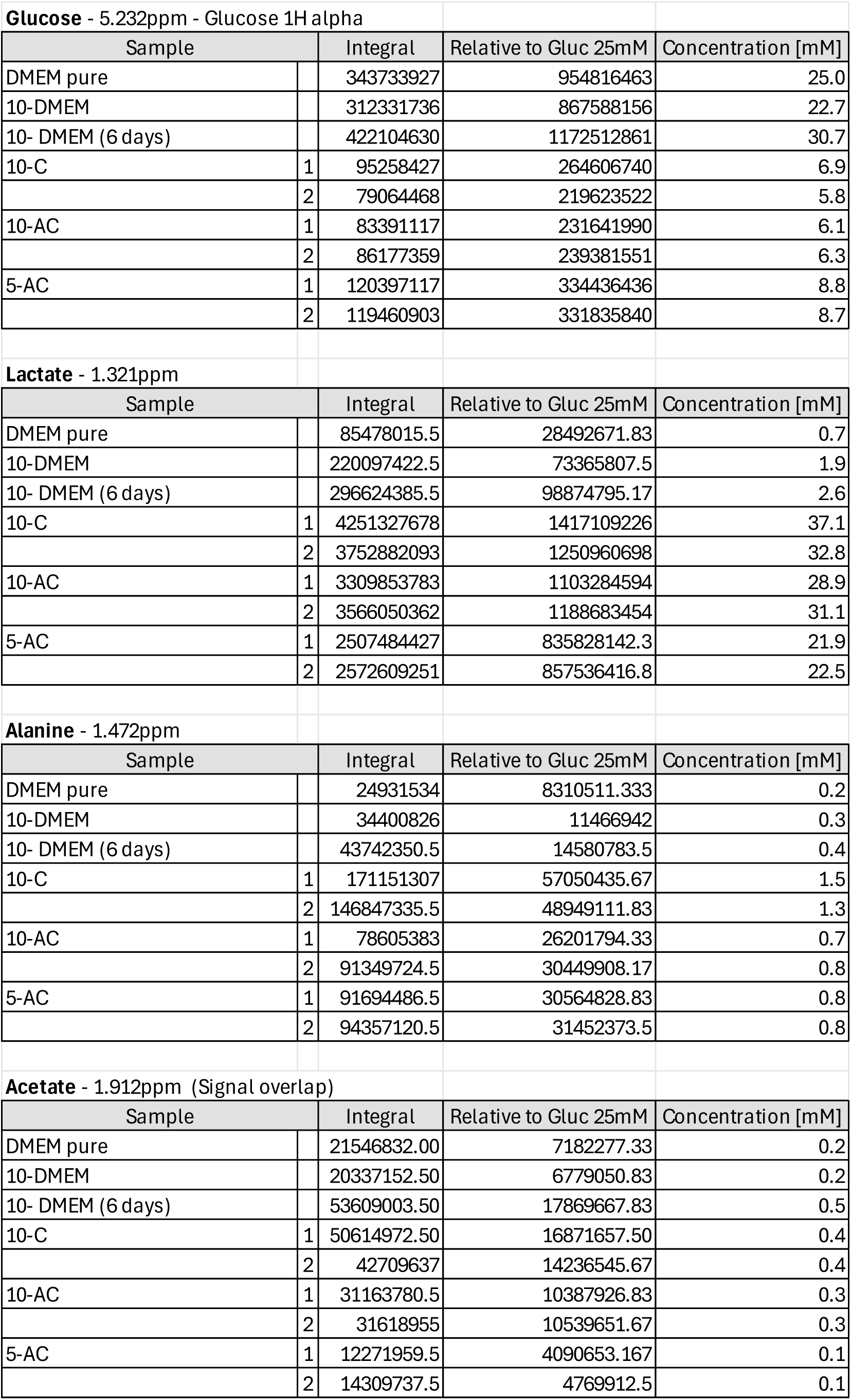
NMR Data and analysis.

**Table ST3:**
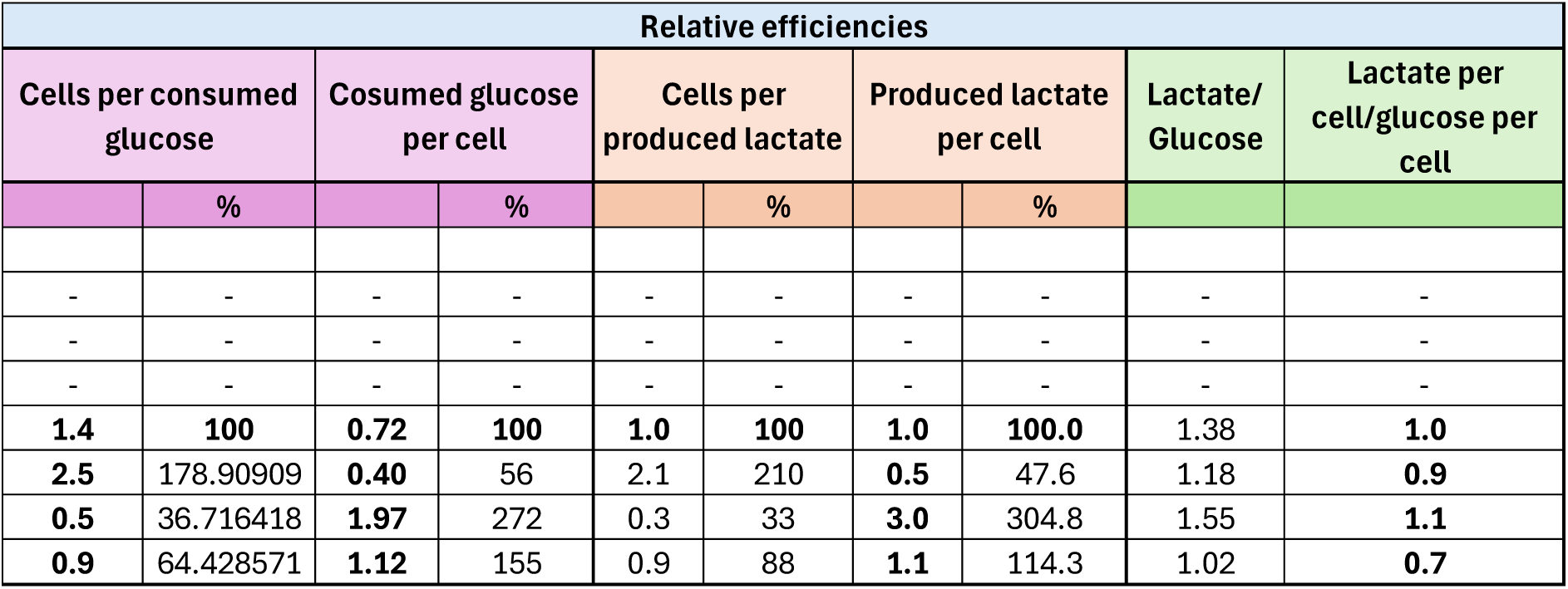
Relative efficiencies in comparison to 10-C samples as standard procedure.

**Table ST4:**
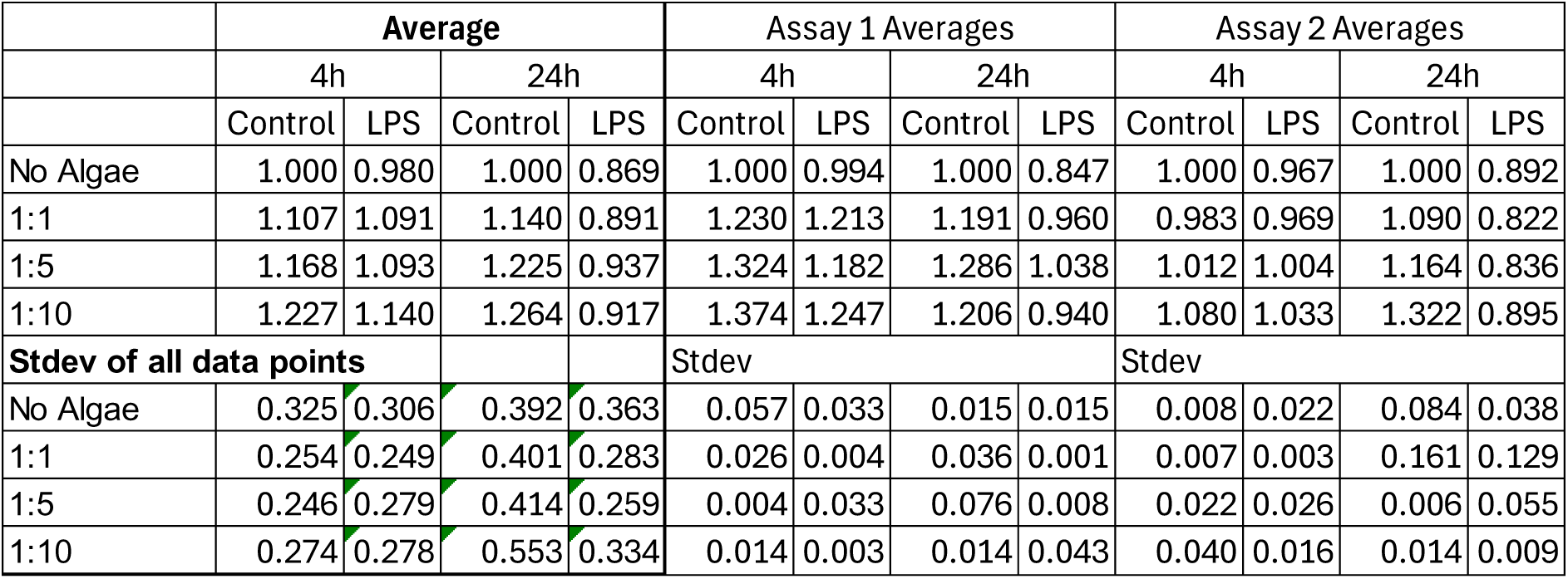
MTT assay data relative to *No Algae -Control* from two independent assays with 2 technical replicates each.

**Table ST5:**
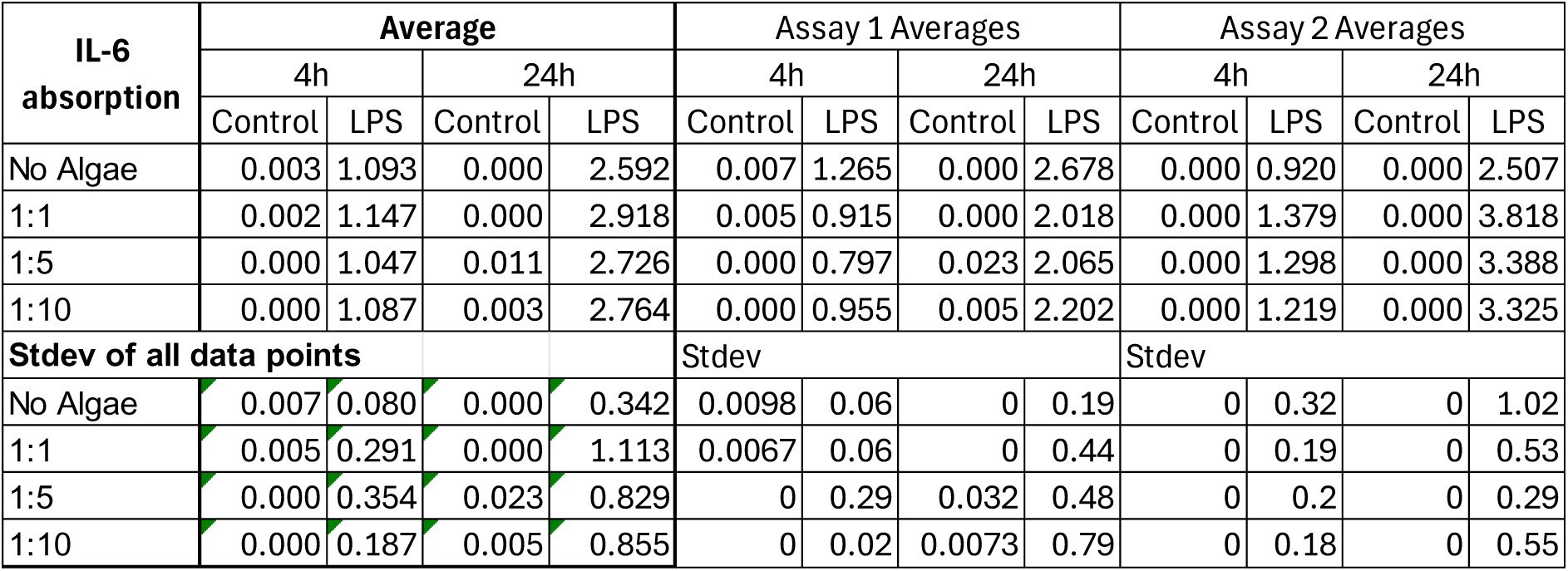
IL-6 absorption from two independent assays with two technical replicates each.

**Figure S3:**
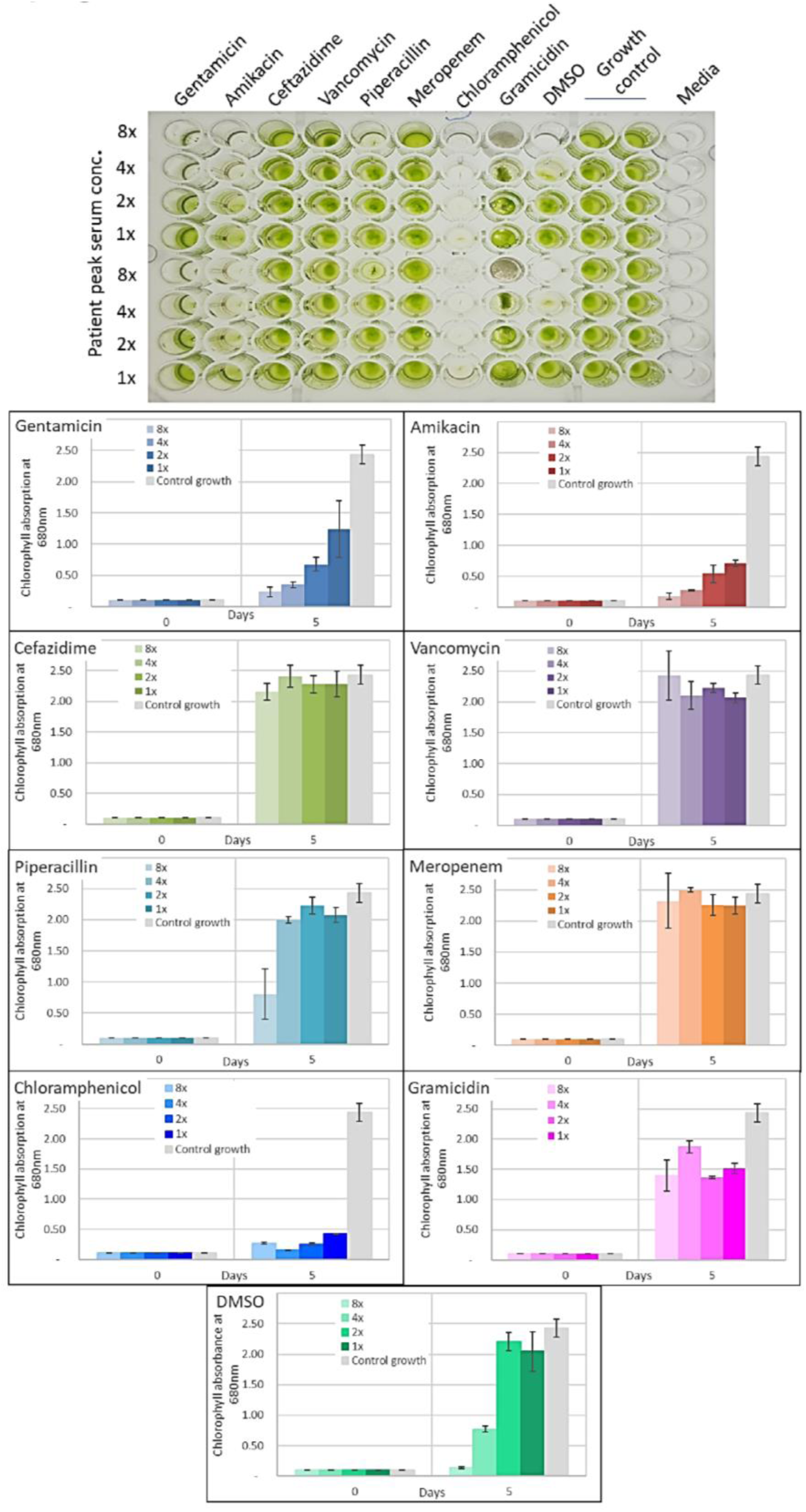
*C. BDH-1* grown in clinically relevant antibiotics at 1×, 2×, 4× and 8× maximum patient serum concentrations according to MIMS (Monthly Index of Medical Specialities) pharmaceutical database, growth plate visual and graphic evaluation (mean±SD, n=2 replicates).

## REFERENCES

1. Verma A, Verma M, Singh A. Chapter 14 - Animal tissue culture principles and applications. In: Verma AS, Singh A, editors. Animal Biotechnology (Second Edition). Boston: Academic Press; 2020. p. 269.

2. Frese L, Darwiche SE, Gunning ME, Hoerstrup SP, von Rechenberg B, Giovanoli P, Calcagni M. Optimizing large-scale autologous human keratinocyte sheets for major burns-Toward an animal-free production and a more accessible clinical application. Health Sci Rep. 2022;5(1):e449.

3. Choi WH, Bae DH, Yoo J. Current status and prospects of organoid-based regenerative medicine. BMB Rep. 2023;56(1):10.

4. Duarte AC, Costa EC, Filipe HAL, Saraiva SM, Jacinto T, Miguel SP, Ribeiro MP, Coutinho P. Animal-derived products in science and current alternatives. Biomater Adv. 2023;151:213428.

5. Subbiahanadar Chelladurai K, Selvan Christyraj JD, Rajagopalan K, Yesudhason BV, Venkatachalam S, Mohan M, Chellathurai Vasantha N, Selvan Christyraj JRS. Alternative to FBS in animal cell culture - An overview and future perspective. Heliyon. 2021;7(8):e07686.

6. Bloch K, Papismedov E, Yavriyants K, Vorobeychik M, Beer S, Vardi P. Immobilized microalgal cells as an oxygen supply system for encapsulated pancreatic islets: a feasibility study. Artif Organs. 2006;30(9):715.

7. Bloch K, Papismedov E, Yavriyants K, Vorobeychik M, Beer S, Vardi P. Photosynthetic oxygen generator for bioartificial pancreas. Tissue Eng. 2006;12(2):337.

8. Haraguchi Y, Kagawa Y, Sakaguchi K, Matsuura K, Shimizu T, Okano T. Thicker three-dimensional tissue from a “symbiotic recycling system” combining mammalian cells and algae. Sci Rep-Uk. 2017;7:41594.

9. Gao B, Hong J, Chen J, Zhang H, Hu R, Zhang C. The growth, lipid accumulation and adaptation mechanism in response to variation of temperature and nitrogen supply in psychrotrophic filamentous microalga Xanthonema hormidioides (Xanthophyceae). Biotechnol Biofuels Bioprod. 2023;16(1):12.

10. Barten RJP, Wijffels RH, Barbosa MJ. Bioprospecting and characterization of temperature tolerant microalgae from Bonaire. Algal Res. 2020;50:102008.

11. Reitman ML. Of mice and men - environmental temperature, body temperature, and treatment of obesity. FEBS Lett. 2018;592(12):2098.

12. van den Tillaart SA, Busard MP, Trimbos JB. The use of distilled water in the achievement of local hemostasis during surgery. Gynecol Surg. 2009;6(3):255.

13. Hazeltine B. CHAPTER 8 - Water Supply. In: Hazeltine B, Bull C, editors. Field Guide to Appropriate Technology. San Diego: Academic Press; 2003. p. 731.

14. Perrineau MM, Zelzion E, Gross J, Price DC, Boyd J, Bhattacharya D. Evolution of salt tolerance in a laboratory reared population of Chlamydomonas reinhardtii. Environ Microbiol. 2014;16(6):1755.

15. Barahoei M, Hatamipour MS, Afsharzadeh S. Direct brackish water desalination using Chlorella vulgaris microalgae. Process Saf Environ Prot. 2021;148:237.

16. Al Bazedi G, Ismail MM, Mugwanya M, Sewilam H. Desalination concentrate microalgae cultivation: biomass production and applications. Sustain Water Resour Manag. 2023;9(4):108.

17. Bartley ML, Boeing WJ, Corcoran AA, Holguin FO, Schaub T. Effects of salinity on growth and lipid accumulation of biofuel microalga Nannochloropsis salina and invading organisms. Biomass Bioenergy. 2013;54:83.

18. Zafar AM, Javed MA, Aly Hassan A, Mehmood K, Sahle-Demessie E. Recent updates on ions and nutrients uptake by halotolerant freshwater and marine microalgae in conditions of high salinity. J Water Process Eng. 2021;44:102382.

19. Wolf J, Ross IL, Radzun KA, Jakob G, Stephens E, Hankamer B. High-throughput screen for high performance microalgae strain selection and integrated media design. Algal Res. 2015;11:313.

20. Whittingham CP, Bermingham M, Hiller RG. The photometabolism of glucose in Chlorella. Queen mary coll london (england); 1963.

21. Savage VM, Allen AP, Brown JH, Gillooly JF, Herman AB, Woodruff WH, West GB. Scaling of number, size, and metabolic rate of cells with body size in mammals. Proc Natl Acad Sci. 2007;104(11):4718.

22. Shaikh S, Lee E, Ahmad K, Ahmad SS, Chun H, Lim J, Lee Y, Choi I. Cell Types Used for Cultured Meat Production and the Importance of Myokines. Foods. 2021;10(10).

23. Harris EH, editor The Chlamydomonas Sourcebook: A Comprehensive Guide to Biology and Laboratory Use 1989.

24. Oey M, Ross IL, Stephens E, Steinbeck J, Wolf J, Radzun KA. RNAi knock-down of LHCBM1, 2 and 3 increases photosynthetic H2 production efficiency of the green alga Chlamydomonas reinhardtii. PLoS One. 2013;8.

25. Guillard RR, Ryther JH. Studies of marine planktonic diatoms. I. Cyclotella nana Hustedt, and Detonula confervacea (cleve) Gran. Can J Microbiol. 1962;8:229.

26. Elisabeth B, Rayen F, Behnam T. Microalgae culture quality indicators: a review. Crit Rev Biotechnol. 2021;41(4):457.

27. Josephine A, Kumar TS, Surendran B, Rajakumar S, Kirubagaran R, Dharani G. Evaluating the effect of various environmental factors on the growth of the marine microalgae, Chlorella vulgaris. Front Mar Sci. 2022;9.

28. Chai S, Shi J, Huang T, Guo Y, Wei J, Guo M, Li L, Dou S, Liu L, Liu G. Characterization of Chlorella sorokiniana growth properties in monosaccharide-supplemented batch culture. PLoS One. 2018;13(7):e0199873.

29. Zhang Z, Schwartz S, Wagner L, Miller W. A greedy algorithm for aligning DNA sequences. J Comput Biol. 2000;7(1-2):203.

30. Stiller JW, McClanahan A. Phyto-specific 16S rDNA PCR primers for recovering algal and plant sequences from mixed samples. Mol Ecol Notes. 2005;5(1):1.

31. Sharma K, Fizet KJ, Montgomery KR, Smeltzer NA, Sikorski MH, Brown KG, Beyke BJ, Burkhart RC, Lynn AN, Grandinetti G. A simple colorimetric experiment using mammalian cell culture to study metabolism. Biochem Mol Biol Educ. 2021;49(2):271.

32. Morgan A, Babu D, Reiz B, Whittal R, Suh LYK, Siraki AG. Caution for the routine use of phenol red – It is more than just a pH indicator. Chem Biol Interact. 2019;310:108739.

33. Ren H, Zhang D. Lactylation constrains OXPHOS under hypoxia. Cell Res. 2024;34(2):91.

34. Kato Y, Inabe K, Haraguchi Y, Shimizu T, Kondo A, Hasunuma T. l-Lactate treatment by photosynthetic cyanobacteria expressing heterogeneous l-lactate dehydrogenase. Sci Rep. 2023;13(1):7249.

35. Vander Heiden MG, Cantley LC, Thompson CB. Understanding the Warburg effect: the metabolic requirements of cell proliferation. Science. 2009;324(5930):1029.

36. Prochownik E, Wang H. The Metabolic Fates of Pyruvate in Normal and Neoplastic Cells. Cells. 2021;10:762.

37. Li Y, Zhao T, Sun W, Gao R, Ma G. Supplementation of alanine improves biomass accumulation and lipid production of Chlorella pyrenoidosa by increasing the respiratory and metabolic processes. J Oceanol Limnol. 2024;42(2):570.

38. Sahuri-Arisoylu M, Mould RR, Shinjyo N, Bligh SWA, Nunn AVW, Guy GW, Thomas EL, Bell JD. Acetate Induces Growth Arrest in Colon Cancer Cells Through Modulation of Mitochondrial Function. Front Nutr. 2021;8:588466.

39. Liu X, Cooper DE, Cluntun AA, Warmoes MO, Zhao S, Reid MA, Liu J, Lund PJ, Lopes M, Garcia BA, et al. Acetate Production from Glucose and Coupling to Mitochondrial Metabolism in Mammals. Cell. 2018;175(2):502–13.e13.

40. Pandian N, Chaurasia R, Chatterjee S, Biswas B, Patra P, Tiwari A, Mukherjee M. Roadmap of algal autotrophic tissue engineering in the avenue of regenerative wound therapy. Mater Adv. 2024;5(19):7516.

41. Alvarez M, Reynaert N, Chávez MN, Aedo G, Araya F, Hopfner U, Fernández J, Allende ML, Egaña JT. Generation of Viable Plant-Vertebrate Chimeras. PLoS One. 2015;10(6):e0130295.

42. Schenck TL, Hopfner U, Chávez MN, Machens HG, Somlai-Schweiger I, Giunta RE, Bohne AV, Nickelsen J, Allende ML, Egaña JT. Photosynthetic biomaterials: a pathway towards autotrophic tissue engineering. Acta Biomater. 2015;15:39.

43. Chávez MN, Schenck TL, Hopfner U, Centeno-Cerdas C, Somlai-Schweiger I, Schwarz C, Machens H-G, Heikenwalder M, Bono MR, Allende ML, et al. Towards autotrophic tissue engineering: Photosynthetic gene therapy for regeneration. Biomaterials. 2016;75:25.

44. Cavalier-Smith T. Chloroplast Evolution: Secondary Symbiogenesis and Multiple Losses. Curr Biol. 2002;12(2):R62.

45. Ma J, Fang Y, Yu H, Yi J, Ma Y, Lei P, Yang Q, Jin L, Wu W, Li H, et al. Recent Advances in Living Algae Seeding Wound Dressing: Focusing on Diabetic Chronic Wound Healing. Adv Funct Mater. 2024;34(2):2308387.

46. McIlroy SE, terHorst CP, Teece M, Coffroth MA. Nutrient dynamics in coral symbiosis depend on both the relative and absolute abundance of Symbiodiniaceae species. Microbiome. 2022;10(1):192.

47. Thomson D, Henry R. Single-step protocol for preparation of plant tissue for analysis by PCR. BioTechniques. 1995;19(3):394, 400.

48. Xu M, McCanna DJ, Sivak JG. Use of the viability reagent PrestoBlue in comparison with alamarBlue and MTT to assess the viability of human corneal epithelial cells. J Pharmacol Toxicol Methods. 2015;71:1.

49. Hotelling H. Analysis of a complex of statistical variables into principal components. J Educ Psychol. 1933;24(6):417.

50. Cloarec O, Dumas ME, Trygg J, Craig A, Barton RH, Lindon JC, Nicholson JK, Holmes E. Evaluation of the orthogonal projection on latent structure model limitations caused by chemical shift variability and improved visualization of biomarker changes in 1H NMR spectroscopic metabonomic studies. Anal Chem. 2005;77(2):517.

