## Supplementary Data for "New glucose-abstaining *Chlorella* algae doubles mammalian cell culture longevity, boosts performance and drops serum needs enabling scaled applications"

Table ST1: Independent experiments (Exp) for resazurine fluorescence data from varying mammalian : algae cell co-cultivation ratios (mean±SD, n=3).

| **Day 2** | | | | | | |
|---|---|---|---|---|---|---|
| | No Algae | 1 : 0.02 | 1 : 0.2 | 1 : 2 | 1 : 20 | 1 : 200 |
| Exp1 | 1.000 | | | | 1.132 | |
| Exp2 | 1.000 | | 0.963 | 1.677 | 3.654 | 1.982 |
| Exp3 | 1.000 | 0.986 | 1.110 | 1.134 | 1.236 | 0.837 |
| Exp4 | 1.000 | | | | 1.039 | |
| Exp5 | 1.000 | | 1.159 | 1.205 | 1.091 | 0.970 |
| Exp6 | 1.000 | 1.272 | 1.346 | 2.111 | 3.159 | 2.224 |
| **Overall Average** | **1.000** | **0.986** | **1.077** | **1.339** | **1.885** | **1.263** |
| **Overall Std. Dev.** | **0.115** | **0.139** | **0.180** | **0.117** | **0.076** | **0.120** |
| **Day 6 (normalized to Day 2 *No Algae* )** | | | | | | |
| Exp1 | 0.715 | | | | 1.050 | |
| Exp3 | 2.246 | 2.256 | 2.322 | 2.365 | 2.604 | 1.746 |
| Exp4 | 0.546 | | | | 1.041 | |
| Exp6 | 2.339 | 2.250 | 2.437 | 2.765 | 5.756 | 2.172 |
| **Overall Average** | **1.462** | **2.253** | **2.379** | **2.565** | **2.613** | **1.959** |
| **Overall Std. Dev.** | **0.207** | **0.029** | **0.182** | **0.103** | **0.047** | **0.046** |

Table ST2: Independent experiments (Exp) to obtain relative fluorescence data normalized to control (10% FBS, 10-C) at day 2 from C2C12 cultivation with or without microalgae in 10% or 5% FBS, respectively (mean±SD, n=3).

| **Day 2** | | | | |
|---|---|---|---|---|
| | **10% FBS (10-C)** | **Algae+10% FBS (10-AC)** | **5% FBS (5-C)** | **Algae+5% FBS (5-AC)** |
| Exp1 | 1 | 1.132 | 0.917 | 0.986 |
| Exp2 | 1 | 3.654 | | |
| Exp3 | 1 | 1.236 | | |
| Exp4 | 1 | 1.039 | 0.685 | 0.837 |
| Exp5 | 1 | 1.084 | | |
| Exp6 | 1 | 3.159 | | |
| **Overall Average** | **1.000** | **1.884** | **0.801** | **0.912** |
| **Overal Std.Dev.** | **0.115** | **0.076** | **0.014** | **0.113** |
| **Day 6 (normalized to Day 2 *10% FBS* )** | | | | |
| Exp1 | 0.715 | 1.050 | 0.521 | 0.837 |
| Exp3 | 2.246 | 2.604 | | |
| Exp4 | 0.546 | 1.041 | 0.433 | 0.709 |
| Exp6 | 2.339 | 5.756 | | |
| **Overall Average** | **1.462** | **2.613** | **0.477** | **0.773** |
| **Overal Std.Dev.** | **0.207** | **0.047** | **0.040** | **0.077** |

Figure S1: Brightfield images of C2C12 cells at 4× magnification on day 0, day 2, and day 6 cultivated with 5% or 10% FBS with or without microalgae.

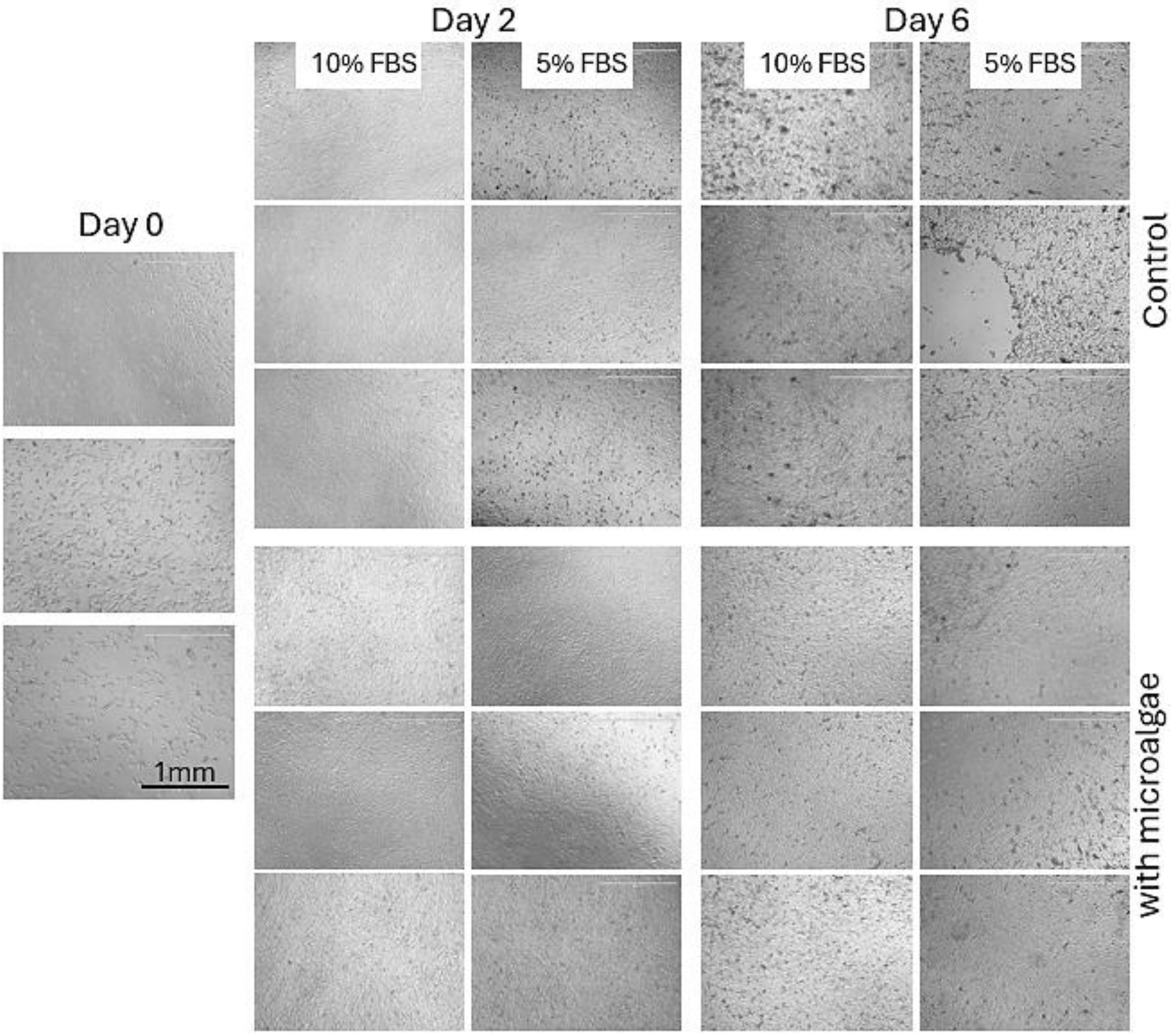

Figure S2: NMR spectra of DMEM pure and complete, as well as pure FBS.

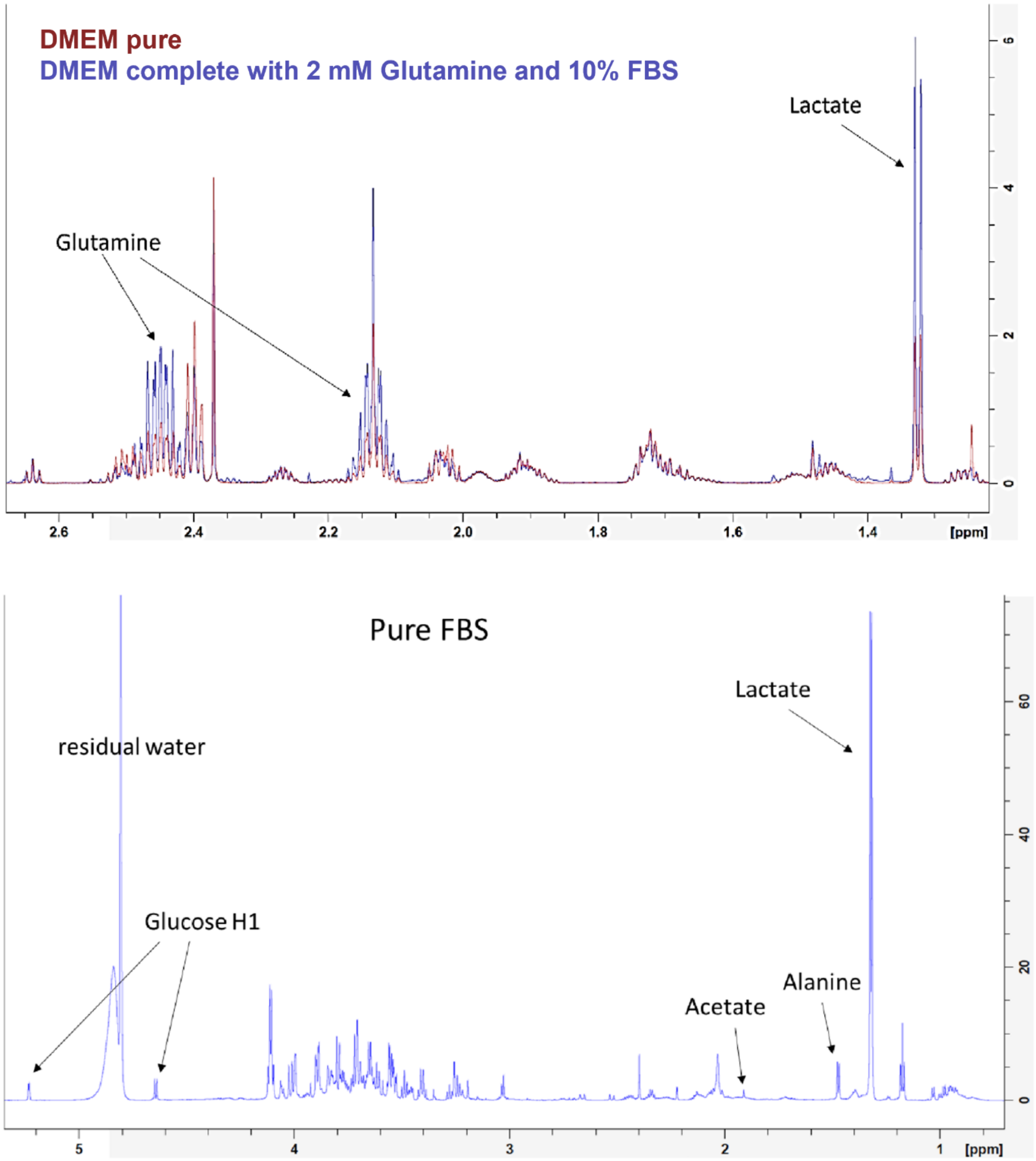

### Table ST2: NMR Data and analysis

**Glucose** - 5.232ppm - Glucose 1H alpha

| Sample | | Integral | Relative to Gluc 25mM | Concentration [mM] |
|---|---|---|---|---|
| DMEM pure | | 343733927 | 954816463 | 25.0 |
| 10-DMEM | | 312331736 | 867588156 | 22.7 |
| 10- DMEM (6 days) | | 422104630 | 1172512861 | 30.7 |
| 10-C | 1 | 95258427 | 264606740 | 6.9 |
| | 2 | 79064468 | 219623522 | 5.8 |
| 10-AC | 1 | 83391117 | 231641990 | 6.1 |
| | 2 | 86177359 | 239381551 | 6.3 |
| 5-AC | 1 | 120397117 | 334436436 | 8.8 |
| | 2 | 119460903 | 331835840 | 8.7 |

**Lactate** - 1.321ppm

| Sample | | Integral | Relative to Gluc 25mM | Concentration [mM] |
|---|---|---|---|---|
| DMEM pure | | 85478015.5 | 28492671.83 | 0.7 |
| 10-DMEM | | 220097422.5 | 73365807.5 | 1.9 |
| 10- DMEM (6 days) | | 296624385.5 | 98874795.17 | 2.6 |
| 10-C | 1 | 4251327678 | 1417109226 | 37.1 |
| | 2 | 3752882093 | 1250960698 | 32.8 |
| 10-AC | 1 | 3309853783 | 1103284594 | 28.9 |
| | 2 | 3566050362 | 1188683454 | 31.1 |
| 5-AC | 1 | 2507484427 | 835828142.3 | 21.9 |
| | 2 | 2572609251 | 857536416.8 | 22.5 |

**Alanine** - 1.472ppm

| Sample | | Integral | Relative to Gluc 25mM | Concentration [mM] |
|---|---|---|---|---|
| DMEM pure | | 24931534 | 8310511.333 | 0.2 |
| 10-DMEM | | 34400826 | 11466942 | 0.3 |
| 10- DMEM (6 days) | | 43742350.5 | 14580783.5 | 0.4 |
| 10-C | 1 | 171151307 | 57050435.67 | 1.5 |
| | 2 | 146847335.5 | 48949111.83 | 1.3 |
| 10-AC | 1 | 78605383 | 26201794.33 | 0.7 |
| | 2 | 91349724.5 | 30449908.17 | 0.8 |
| 5-AC | 1 | 91694486.5 | 30564828.83 | 0.8 |
| | 2 | 94357120.5 | 31452373.5 | 0.8 |

**Acetate** - 1.912ppm (Signal overlap)

| Sample | | Integral | Relative to Gluc 25mM | Concentration [mM] |
|---|---|---|---|---|
| DMEM pure | | 21546832.00 | 7182277.33 | 0.2 |
| 10-DMEM | | 20337152.50 | 6779050.83 | 0.2 |
| 10- DMEM (6 days) | | 53609003.50 | 17869667.83 | 0.5 |
| 10-C | 1 | 50614972.50 | 16871657.50 | 0.4 |
| | 2 | 42709637 | 14236545.67 | 0.4 |
| 10-AC | 1 | 31163780.5 | 10387926.83 | 0.3 |
| | 2 | 31618955 | 10539651.67 | 0.3 |
| 5-AC | 1 | 12271959.5 | 4090653.167 | 0.1 |
| | 2 | 14309737.5 | 4769912.5 | 0.1 |

Table ST3: Relative efficiencies in comparison to 10-C samples as standard procedure.

| Relative efficiencies | | | | | | | | | |
|---|---|---|---|---|---|---|---|---|---|
| Cells per consumed glucose | | Cosumed glucose per cell | | Cells per produced lactate | | Produced lactate per cell | | Lactate/ Glucose | Lactate per cell/glucose per cell |
| | % | | % | | % | | % | | |
| - | - | - | - | - | - | - | - | - | - |
| - | - | - | - | - | - | - | - | - | - |
| - | - | - | - | - | - | - | - | - | - |
| **1.4** | **100** | **0.72** | **100** | **1.0** | **100** | **1.0** | **100.0** | 1.38 | **1.0** |
| **2.5** | 178.90909 | **0.40** | 56 | 2.1 | 210 | **0.5** | 47.6 | 1.18 | **0.9** |
| **0.5** | 36.716418 | **1.97** | 272 | 0.3 | 33 | **3.0** | 304.8 | 1.55 | **1.1** |
| **0.9** | 64.428571 | **1.12** | 155 | 0.9 | 88 | **1.1** | 114.3 | 1.02 | **0.7** |

Table ST4: MTT assay data relative to *No Algae -Control* from two independent assays with 2 technical replicates each.

| | **Average** | | | | Assay 1 Averages | | | | Assay 2 Averages | | | |
|---|---|---|---|---|---|---|---|---|---|---|---|---|
| | 4h | | 24h | | 4h | | 24h | | 4h | | 24h | |
| | Control | LPS | Control | LPS | Control | LPS | Control | LPS | Control | LPS | Control | LPS |
| No Algae | 1.000 | 0.980 | 1.000 | 0.869 | 1.000 | 0.994 | 1.000 | 0.847 | 1.000 | 0.967 | 1.000 | 0.892 |
| 1:1 | 1.107 | 1.091 | 1.140 | 0.891 | 1.230 | 1.213 | 1.191 | 0.960 | 0.983 | 0.969 | 1.090 | 0.822 |
| 1:5 | 1.168 | 1.093 | 1.225 | 0.937 | 1.324 | 1.182 | 1.286 | 1.038 | 1.012 | 1.004 | 1.164 | 0.836 |
| 1:10 | 1.227 | 1.140 | 1.264 | 0.917 | 1.374 | 1.247 | 1.206 | 0.940 | 1.080 | 1.033 | 1.322 | 0.895 |
| **Stdev of all data points** | | | | | Stdev | | | | Stdev | | | |
| No Algae | 0.325 | 0.306 | 0.392 | 0.363 | 0.057 | 0.033 | 0.015 | 0.015 | 0.008 | 0.022 | 0.084 | 0.038 |
| 1:1 | 0.254 | 0.249 | 0.401 | 0.283 | 0.026 | 0.004 | 0.036 | 0.001 | 0.007 | 0.003 | 0.161 | 0.129 |
| 1:5 | 0.246 | 0.279 | 0.414 | 0.259 | 0.004 | 0.033 | 0.076 | 0.008 | 0.022 | 0.026 | 0.006 | 0.055 |
| 1:10 | 0.274 | 0.278 | 0.553 | 0.334 | 0.014 | 0.003 | 0.014 | 0.043 | 0.040 | 0.016 | 0.014 | 0.009 |

Table ST5: IL-6 absorption from two independent assays with two technical replicates each.

| **IL-6 absorption** | **Average** | | | | Assay 1 Averages | | | | Assay 2 Averages | | | |
|---|---|---|---|---|---|---|---|---|---|---|---|---|
| | 4h | | 24h | | 4h | | 24h | | 4h | | 24h | |
| | Control | LPS | Control | LPS | Control | LPS | Control | LPS | Control | LPS | Control | LPS |
| No Algae | 0.003 | 1.093 | 0.000 | 2.592 | 0.007 | 1.265 | 0.000 | 2.678 | 0.000 | 0.920 | 0.000 | 2.507 |
| 1:1 | 0.002 | 1.147 | 0.000 | 2.918 | 0.005 | 0.915 | 0.000 | 2.018 | 0.000 | 1.379 | 0.000 | 3.818 |
| 1:5 | 0.000 | 1.047 | 0.011 | 2.726 | 0.000 | 0.797 | 0.023 | 2.065 | 0.000 | 1.298 | 0.000 | 3.388 |
| 1:10 | 0.000 | 1.087 | 0.003 | 2.764 | 0.000 | 0.955 | 0.005 | 2.202 | 0.000 | 1.219 | 0.000 | 3.325 |
| **Stdev of all data points** | | | | | Stdev | | | | Stdev | | | |
| No Algae | 0.007 | 0.080 | 0.000 | 0.342 | 0.0098 | 0.06 | 0 | 0.19 | 0 | 0.32 | 0 | 1.02 |
| 1:1 | 0.005 | 0.291 | 0.000 | 1.113 | 0.0067 | 0.06 | 0 | 0.44 | 0 | 0.19 | 0 | 0.53 |
| 1:5 | 0.000 | 0.354 | 0.023 | 0.829 | 0 | 0.29 | 0.032 | 0.48 | 0 | 0.2 | 0 | 0.29 |
| 1:10 | 0.000 | 0.187 | 0.005 | 0.855 | 0 | 0.02 | 0.0073 | 0.79 | 0 | 0.18 | 0 | 0.55 |

Figure S3*: C. BDH-1* grown in clinically relevant antibiotics at 1×, 2×, 4× and 8× maximum patient serum concentrations according to MIMS (Monthly Index of Medical Specialities) pharmaceutical database, growth plate visual and graphic evaluation (mean±SD, n=2 replicates).

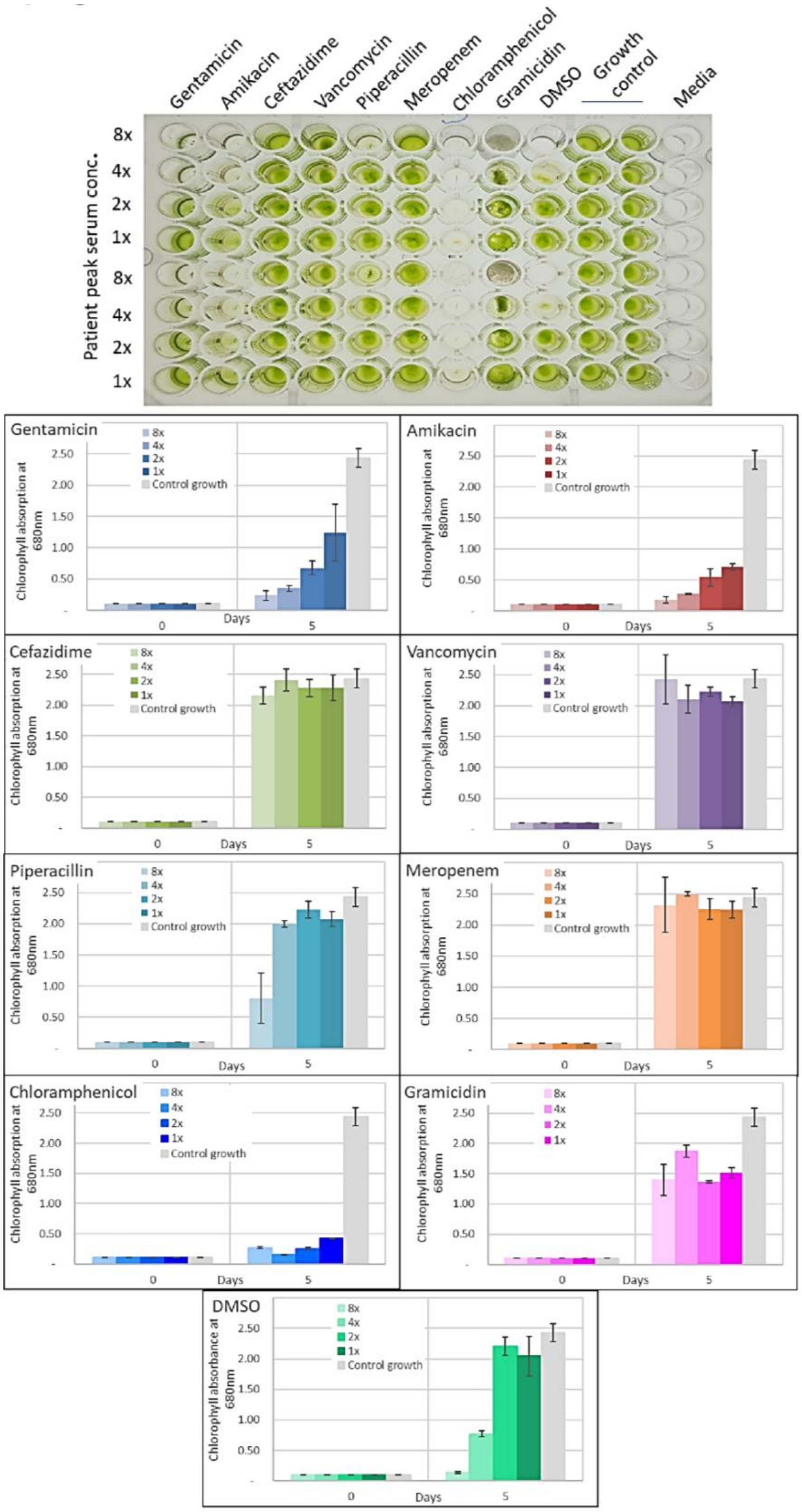
